# Exploratory Factor Analysis with Structured Residuals for Brain Imaging Data

**DOI:** 10.1101/2020.02.06.933689

**Authors:** Erik-Jan van Kesteren, Rogier A. Kievit

## Abstract

Dimension reduction is widely used and often necessary to reduce high dimensional data to a small number of underlying variables, making subsequent analyses and their interpretation tractable. One popular technique is Exploratory Factor Analysis (EFA), used by cognitive neuroscientists to reduce measurements from a large number of brain regions to a tractable number of factors. However, dimension reduction often ignores relevant a priori knowledge about the structure of the data. For example, it is well established that the brain is highly symmetric. In this paper, we (a) show the adverse consequences of ignoring a priori structure in factor analysis, (b) propose a technique to accommodate structure in EFA using structured residuals (EFAST), and (c) apply this technique to three large and varied brain imaging datasets, demonstrating the superior fit and interpretability of our approach. We provide an R software package to enable researchers to apply EFAST to other suitable datasets.

## 1 Introduction

Scientists in vastly different disciplines often face a similar problem: the challenges of dimensionality. Data collection and acquisition may yield far more variables than can be tractably analyzed, yet omitting large proportions of the data is equally undesirable. In the fields of statistics and mathematics, there have been numerous developments that deal with these challenges by means of *dimension reduction*. In such approaches, researchers take a high dimensional dataset and reduce it to a (much) smaller number of dimensions, often called factors or components, for further analysis.

A particularly popular dimension reduction technique is Exploratory Factor Analysis (EFA). A precursor of modern EFA was invented by Spearman (Bartholomew, 1995), who developed it to reduce performance scores on a large battery of cognitive ability tests into one, or a small number, of ability factors. EFA models the observed covariance matrix of a set of *P* variables by assuming there are *M < P* factors, which predict the values on the observed variables. For example, an underlying fluid intelligence factor may partially predict scores on a matrix reasoning test (Horn and Cattell, 1967).

Many data reduction techniques exist beyond EFA, including principal components analysis (PCA), Partial Least Squares (PLS), Independent Component Analysis (ICA), and many more beyond our current scope (see Roweis and Ghahramani, 1999; Sorzano et al., 2014). All of these techniques aim to approximate the observed data by means of a lower-dimensional representation. These techniques, although powerful, share a particular limitation, at least in their canonical implementations, namely that they cannot easily integrate prior knowledge of (additional) covariance structure present in the data. In other words, all observed covariation is modeled by the underlying factor structure.

Dimension reduction techniques have been used more recently in emerging fields such as cognitive neuroscience. High dimensional individual differences in brain structure or function (e.g., volume or activity measures across dozens of regions or even thousands of voxels) are reduced to a smaller number of factors, which are then used, for instance, to study morphological differences in schizophrenia (Tien et al., 1996), how cortical structure relates to behavioural measures (Colibazzi et al., 2008), and to examine age-related differences in brain structure (de Mooij et al., 2018). However, one key challenge when reducing the dimensionality of such structural (and functional) brain data is that of *symmetry*: Much like other body parts, contralateral (left/right) brain regions are highly correlated due to developmental and genetic mechanisms which govern the gross morphology of the brain. Ignoring this prior information will adversely affect the dimension reduction step, leading to worse representation of the high-dimensional data by the extracted factors. Simple workarounds, such as averaging left and right into a single index per region, have other drawbacks: they throw away information, preclude the discovery of (predominantly) lateralized factors, and prevent the study of (a)symmetry as a topic of interest in and of itself. We believe that many data reduction problems in social, cognitive, and behavioural sciences have a similar structure: residual structure is known, but precise theory about the underlying factor structure is not. (Asparouhov and Muthén, 2009). As such, although we focus on imaging data, our approach is likely more widely applicable.

Other classes of techniques, developed largely within psychometrics, can naturally accommodate such additional structure. These techniques started with multitrait - multimethod (MTMM) matrices (Campbell and Fiske, 1959) and later confirmatory factor analysis (CFA) with residual covariances (e.g., Kenny, 1976). MTMM is designed to extract factors when these factors are measured in different ways: when measuring personality through a self-report questionnaire and behaviour ratings, there are factors that explain correlation among items corresponding to a specific trait such as ‘extraversion’, and there are factors that explain additional correlation between items because they are gathered using the same methods (self-report and behavioural ratings). Thus, MTMM techniques separate the correlation matrix into two distinct, summative parts: correlation due to the traits of central interest, and correlation due to the measurement methods. However, MTMM requires a priori knowledge of the trait structure (e.g., the OCEAN model of personality).

Here we propose a manner in which to instead conduct purely *exploratory* factor analysis (e.g. across many brain regions), whilst incorporating prior structure knowledge (e.g. symmetry). Standard implementations of EFA, CFA, and MTMM are inadequate to estimate factor structure under these circumstances, as they do not simultaneously allow for exploration and the incorporation of residual structure. We improve upon these procedures by developing a method called *Exploratory factor analysis with structured residuals* (EFAST). We show that EFAST outperforms EFA in empirically plausible scenarios, and that ignoring the problem of structured residuals in these scenarios adversely affects inferences.

Note that we are not the first to suggest using structured residuals in EFA to take into account prior knowledge about structure in the observed variables. Adding covariances among residuals is a common method to take into account features of the data-generating process (e.g., Cole et al., 2007), and this has been possible in the context of EFA since the release of the ESEM capability in `MPlus` (Asparouhov and Muthén, 2009) and in `lavaan` (Rosseel, 2019). In the context of neuroscientific data, similar methods in accounting for structure in dimension reduction have been researched by De Munck et al. (2002) in source localization for EEG/MEG. Our goal for this paper is to provide a compelling argument for the use of such structured residuals from the point of view of cognitive neuroscience, as well as a user-friendly, open-source implementation of this method for dimension reduction in real-world datasets.

This paper is structured as follows. First, we explain why using standard EFA or CFA for brain imaging data may lead to undesirable results, and we develop EFAST based on novel techniques from structural equation modeling (SEM). Then, we show that EFAST performs well in simulations, demonstrating superior performance compared to EFA in terms of factor recovery, factor covariance estimation, and the number of extracted factors when dealing with symmetry. Third, we illustrate EFAST in a large neuroimaging cohort (Cam-CAN; Shafto et al., 2014). We illustrate EFAST for three distinct datasets: Grey matter volume, white matter microstructure and within-subject fMRI functional connectivity. We show how EFAST outperforms EFA both conceptually and statistically in all three datasets, showing the generality of our technique. We conclude with an overview and suggestions for further research.

Accompanying this paper, we provide tools for researchers to use and expand upon with their own datasets. These tools take the form of (a) an R package called `efast` and a tutorial with example code (https://github.com/vankesteren/efast), and (b) synthetic data and code to reproduce the empirical examples in Section 4 and the simulations in Section 3 (https://github.com/vankesteren/efast_code).

## 2 Factor analysis with structured residuals

In this section, we compare and contrast existing approaches in their ability to perform factor analysis in an exploratory way while at the same time accounting for residual structure. We discuss new developments in the field of exploratory structural equation modeling (ESEM) that enable simultaneous estimation of exploratory factors and structured residuals, after which we develop the EFAST model as an ESEM with a single exploratory block. We will use brain morphology data with bilateral symmetry as our working example throughout, although the principles here can be generalized to datasets with similar properties.

EFA, as implemented in software programs such as SPSS, R, and Mplus, models the observed correlation matrix through two summative components: the factor loading matrix **Λ**, relating the predefined number *M* of factors to the observed variables, and a diagonal residual variance matrix **Θ**, signifying the variance in the observed variables unexplained by the factors. Using maximum likelihood, principal axis factoring, or least squares (Harman and Jones, 1966), the factor loadings and residual variances are estimated such that the implied correlation matrix **Σ**= **ΛΛ**^*T*^ + **Θ** best approximates the observed correlation matrix ****S****. After estimation, the factor loadings are rotated to their final interpretable solution using objectives such as oblimin, varimax, or geomin (Bernaards and Jennrich, 2005).

We illustrate the challenge and the rationale behind our approach in Figure 1. The true correlation matrix is highlighted on the left, with correlations due to three factors shown as diagonal blocks. However, there is also considerable off-diagonal structure: the secondary diagonals show a symmetry pattern similar to that observed in real-world brain structure data (Taylor et al., 2017). The top panel of the figure shows that a traditional EFA approach will separate this data matrix into two components: (a) covariance due to the hypothesized factor structure and (b) the diagonal residual matrix. The key challenge is that EFA will attempt to approximate all the off-diagonal elements of the correlation matrix through the factors, even if this adversely affects the recovery of the true factor structure. Performing EFA with such a symmetry pattern may affect the factor solution in a variety of ways. For instance, in this toy example, the EFA model requires more than 12 factors to properly represent the data, instead of the three factors specified (see Appendix A). In other words, in such cases it is essential to incorporate the known residual structure via a set of additional assumptions.

**Figure 1:**
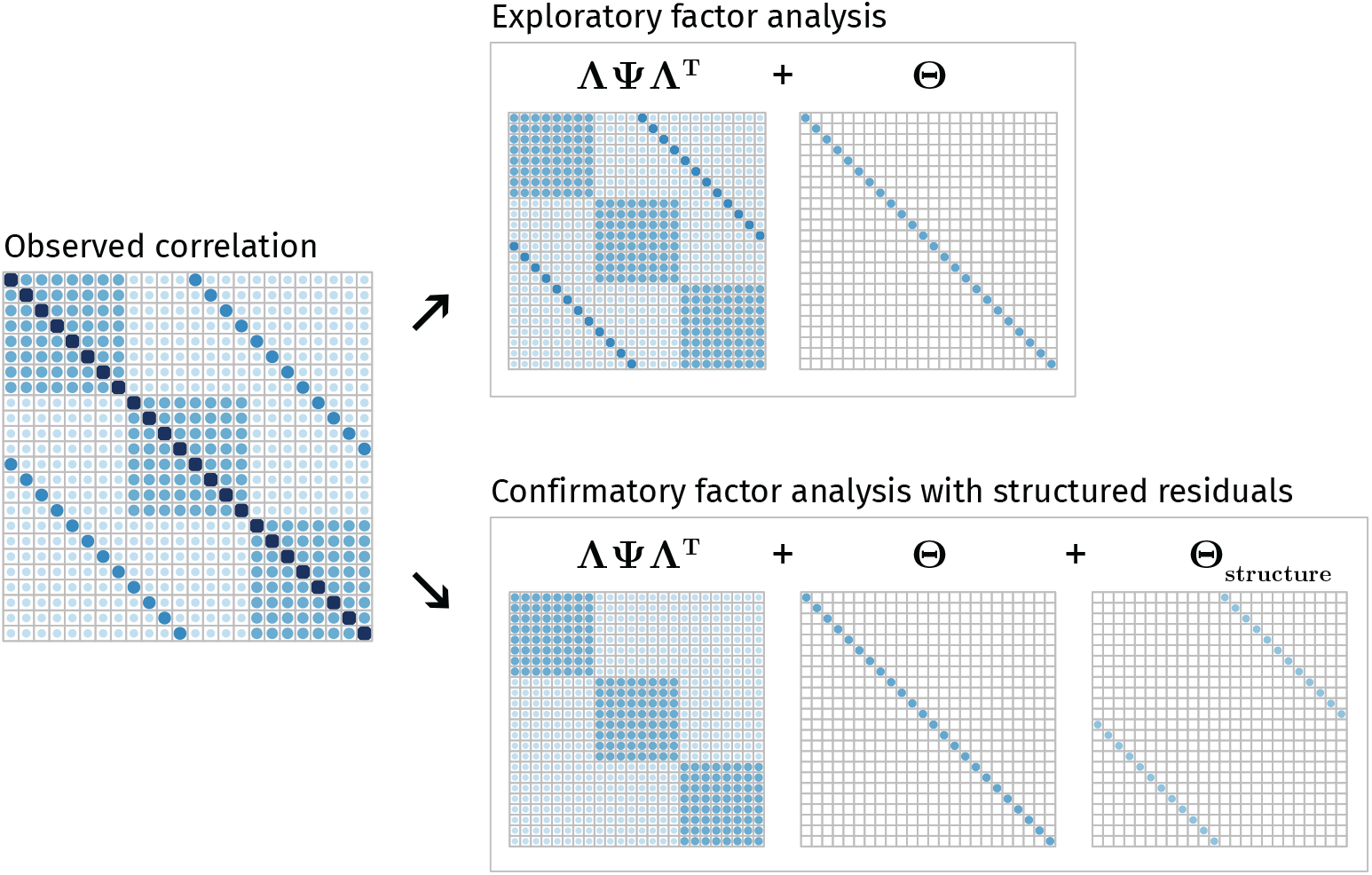
Example observed correlation matrix and its associated decomposition according to EFA (top) and according to CFA (bottom) into a factor-implied correlation component (**ΛΨΛ**^*T*^), residual variance component **Θ**, and – in CFA with residual structure only – residual structure component.

As an alternative to EFA, we may implement a Confirmatory Factor Analysis (CFA) instead. In contrast to EFA, CFA imposes a priori constraints on the **Λ** matrix: some observed variables do not load on some factors. Moreover, in contrast to standard EFA approaches, residual structure can be easily implemented in CFA using standard SEM software such as `lavaan` (Rosseel, 2012). In other words, CFA would allow us to tackle the problem in Figure 1: We can allow for the residual structure known a priori to be present in the data. By allowing for the residual structure in the data, a CFA yields the implied matrices shown in the bottom panel of Figure 1, retrieving the correct factor loadings, residual variance, and residual structure. However, this is only possible because in this toy example we *know* the factor structure - In many empirical situations this is precisely what we wish to discover. In the absence of theory about the underlying factors, it is thus not possible to benefit from these features of CFA.

As such, we need an approach that can combine the strengths of EFA (estimating the factor structure in the absence of strong a priori theory) with those from CFA (the potential to allow for a priori residual structure). Here, we propose a hybrid between the two, which we call *exploratory factor analysis with structured residuals*, or EFAST. In order to implement and estimate these models, we make use of recent developments in the field of structural equation modeling (SEM). In the next section, we explain how these developments make EFAST estimation possible.

### 2.1 Exploratory SEM

Exploratory SEM (ESEM) is an extension to SEM which allows for blocks of exploratory factor analysis within the framework of confirmatory SEM (Jöreskog, 1969; Brown, 2006; Asparouhov and Muthén, 2009; Guàrdia-Olmos et al., 2009; Marsh et al., 2014; Rosseel, 2019). ESEM is a two-step procedure. In the first step, a regular SEM model is estimated, where each of the EFA blocks have a diagonal latent covariance matrix **Ψ** and the **Λ** matrix of each block is of transposed echelon form, meaning all elements above the diagonal are constrained to 0. For a nine-variable, three-factor EFA block *b* the matrices would then be:

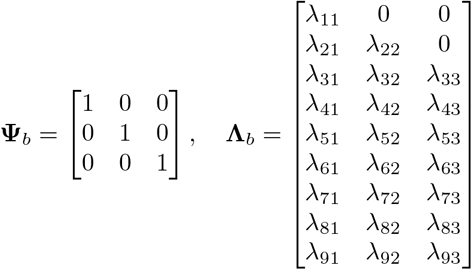

This means there are 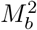 constraints for each EFA block *b*. This is the same number of constraints as conventional EFA (Asparouhov and Muthén, 2009). The second step in ESEM is to rotate the solution using a rotation matrix ****H****. Just as in regular EFA, this rotation matrix is constructed using objectives such as geomin or oblimin. In ESEM, the rotation affects the factor loadings and latent covariances of the EFA blocks, but also almost all other parameters in the model (Asparouhov and Muthén (2009) provide an overview of how rotation changes these parameter estimates). Despite these changes, a key property of ESEM is that different rotation solutions lead to the same overall model fit.

ESEM has long been available only in Mplus (Muthén and Muthén, 1998; Asparouhov and Muthén, 2009). More recently, it has become available in open sourced `R` packages psych (for specific models, Revelle, 2018) as well as `lavaan` (since version 0.6-4, Rosseel, 2019) – a comprehensive package for structural equation modeling. An example of a basic EFA model using `lavaan` syntax with 3 latent variables and 9 observed variables is the following:

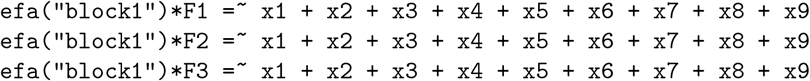

In effect, this model specifies three latent variables (`F1`, `F2`, and `F3`) which are each indicated by all 9 observed variables (`x1` to `x9`). The `efa(“block1”)` part is a modifier for this model which imposes the constraints on **Ψ** and **Λ** mentioned above. For a more detailed explanation of the `lavaan` syntax, see Rosseel (2012). Figure 2 shows a comparison of the factor loadings obtained using conventional factor analysis (`factanal()` in `R`) and `lavaan`’s `efa()` modifier. As shown, the solution obtained is exactly the same, with perfect correlation among the loadings for each of the factors.

**Figure 2:**
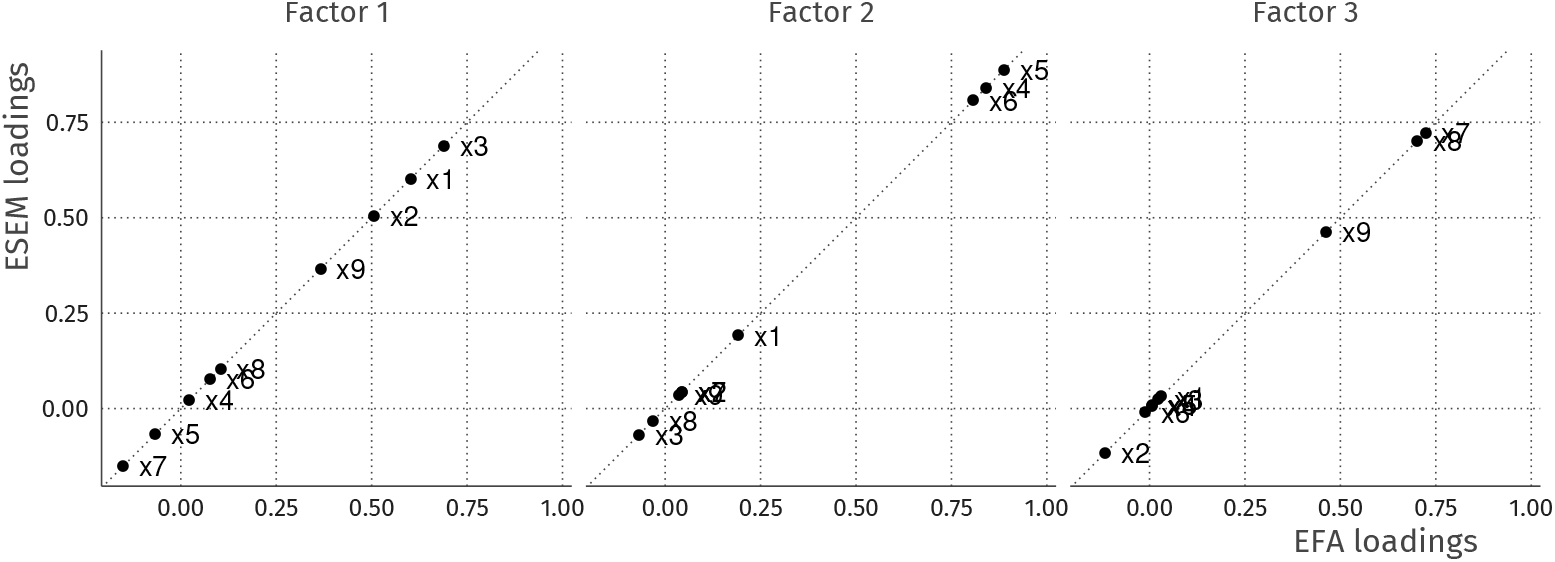
Exploratory factor analysis of 9 variables in the Holzinger and Swineford (1939) dataset. On the y-axis are the estimated factor loadings using the oblimin rotation functionality in `lavaan` version 0.6-4, and the loadings on the x-axis are derived from `factanal` with oblimin rotation from the `GPArotation` package (Bernaards and Jennrich, 2005). The loadings are all on the diagonal with a correlation of 1, meaning the solutions obtained from these different methods are equal.

With this tool as the basis for model estimation, the next section provides a detailed development of the construction of EFAST models.

### 2.2 EFAST models

We propose using EFA with corrections for contralateral covariance within the ESEM framework. The corrections we propose are the same as in MTMM models or CFA with residual covariance. In EFAST the method factors use CFA, and the remaining correlations are explained by EFA. Thus, unlike standard MTMM methods, EFAST contains *exploratory* factor analysis on the trait side, as the factor structure of the traits is unknown beforehand: the goal of the analysis is to extract an underlying low-dimensional set of features which explain the observed correlations as well as possible. For our running example of brain imaging data with contralateral symmetry, we consider each ROI a “method” factor, loading on only two regions. Note that in the context of brain imaging, Lövdén et al. (2013, Figure 1, model A) have had similar ideas, but their factor analysis operates on the level of left-right combined ROIs rather than individual ROIs.

The EFAST model has *M* exploratory factors in a single EFA block, and one method factor per homologous ROI pair, each with loadings constrained to 1 and its own variance estimated. The estimated variance of the method factors then represents the amount of covariance due to symmetry – over and above the covariance represented by the traits. In Figure 3, the model is displayed graphically for a simplified example with 6 ROIs in each hemisphere.

**Figure 3:**
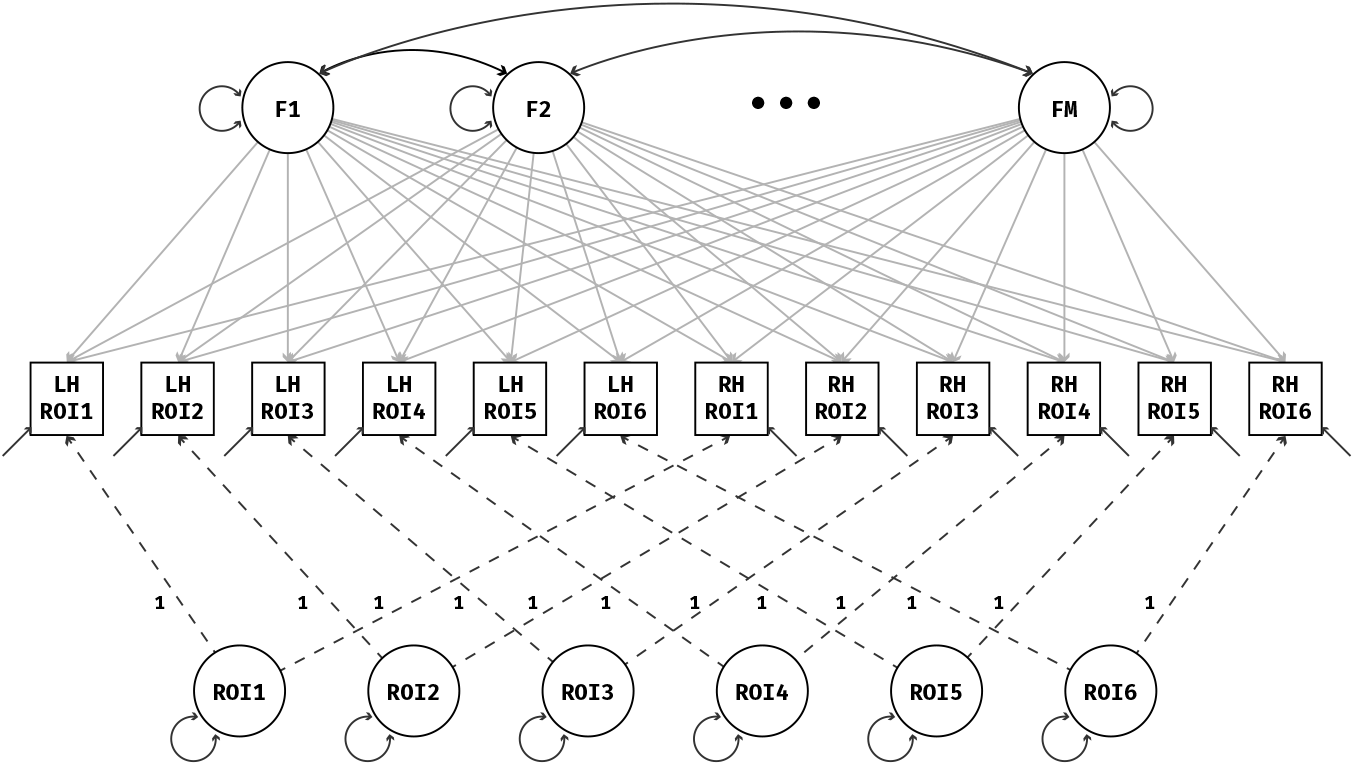
EFAST model with morphology of 6 regions of interest measured in the left hemisphere (LH) and right hemisphere (RH). The dashed lines indicate fixed loadings, the two-headed arrows indicate variance/covariance parameters. The method factors are constrained to be orthogonal, and the loadings of the *M* traits are estimated in an exploratory way.

An alternative parametrization for this model is also available. Specifically, we can use the correlations between the residuals of the observed variables instead of method factors with freely estimated variances. In the SEM framework, this would amount to moving the symmetry structure from the factor-explained matrix (**ΛΨΛ**^*T*^) to the residual covariance matrix **Θ**. This model is exactly equivalent, meaning the same correlation matrix decomposition, the same factor structure, and the same model fit will be obtained. However, we favour the method factor parametrization as it is closer to MTMM-style models, it is easier to extract potentially relevant metrics such as a ‘lateralisation coefficient’, and easier to extend to other data situations where multiple indicators load on each method factor.

To implement the EFAST model we use the package `lavaan`, which allows for easy scaling of the input data, different estimation methods, missing data handling through full information maximum likelihood, and more. Estimation of the model in Figure 3 can be done with a variety of methods. Here we use the default maximum likelihood estimation method as implemented in `lavaan`. Accompanying this paper, we are making available a convenient `R` package called `efast` that can fit EFAST models for datasets with residual structure due to symmetry. For more implementation details, the package and its documentation can be found at https://github.com/vankesteren/efast.

In the next section, we show how our implementation of EFAST compares to regular EFA in terms of factor loading estimation, factor covariance estimation, as well as the estimated number of factors.

## 3 Simulations

In this section, we use simulated data to examine different properties of EFAST models when compared to regular EFA in controlled conditions. The purpose of this simulation is not an exhaustive investigation, but rather a pragmatically focused study of data properties (neuro)scientists wishing to use this technique are likely to encounter. First, we explain how data were simulated to follow a specific correlation structure, approximating observed empirical data such as that in the Cam-CAN study (Section 4, Figure 10). Then, we investigate the effects of structured residuals on the extracted factors from EFA and EFAST: in several different conditions, we investigate how the estimation of factor loadings, the covariances between factors, and the number of factors changes with increasing symmetry.

### 3.1 Data generation

Data were generated following a controlled population correlation matrix **Σ**_*true*_. This matrix represents the true correlation between measurements of brain structure in 17 left-hemisphere and 17 right-hemisphere regions of interest. An example correlation matrix from our data-generating mechanism is shown in Figure 4.

**Figure 4:**
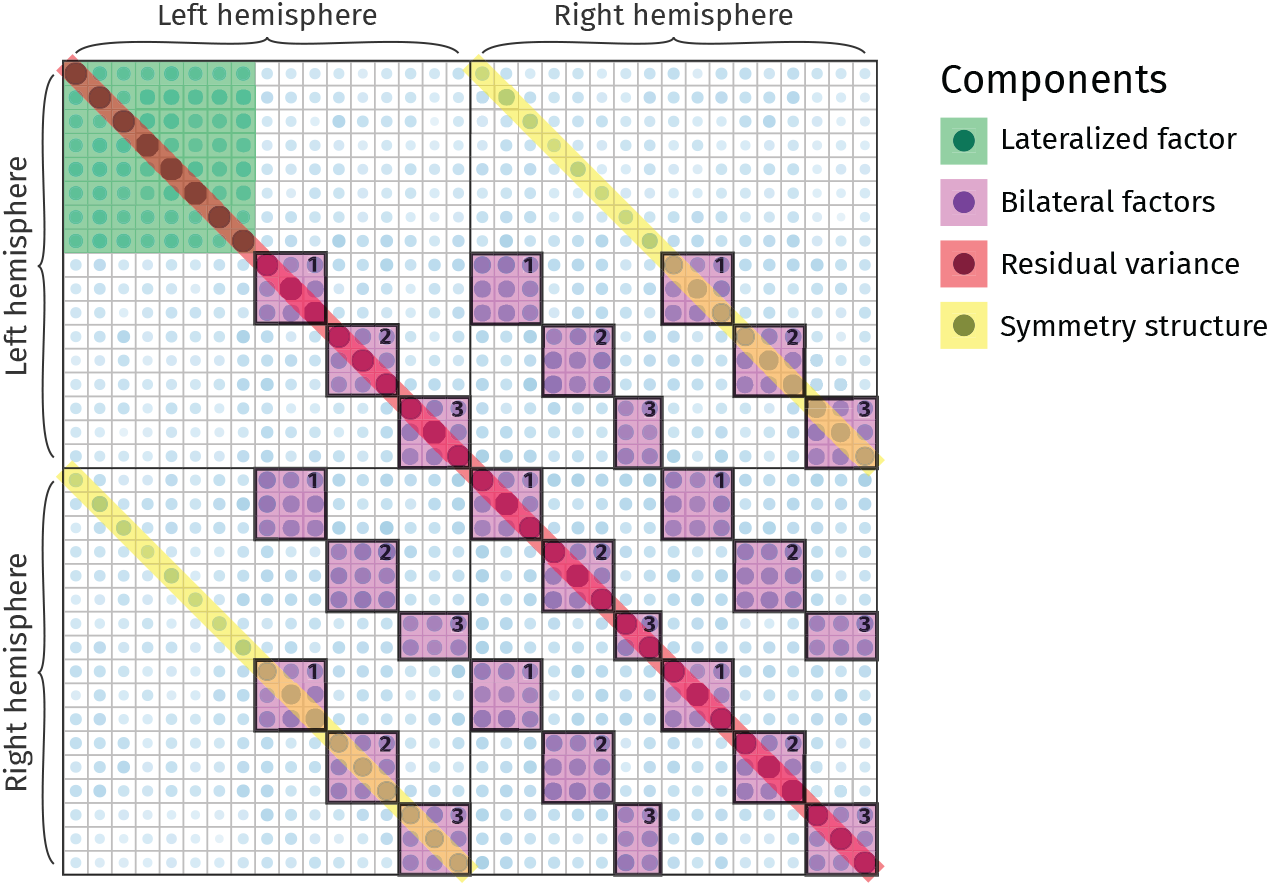
Example covariance matrix of the data-generating mechanism used in the simulations. This matrix results from simulated data of 650 brain images, with a factor loading of .595 for the lateralized factor, a loading of .7 for the remaining factors, a factor correlation of .5, and a symmetry correlation of .2. The first 17 variables indicate regions of interest (ROIs) in the left hemisphere, and the remaining variables indicate their contralateral homologues. Note the secondary diagonals, indicating contralateral symmetry, and the block of 8 variables in the top left resulting from the lateralized factor.

**Σ**_*true*_ was constructed through the summation of three separate matrices, as in the lower panel of Figure 1:

1. The factor component **Σ**_*factor*_ is constructed as **ΛΨΛ**^*T*^, where the underlying factor covariance matrix **Ψ** can be either an identity matrix (orthogonal factors) or a matrix with nonzero off-diagonal elements (oblique factors). There are four true underlying factors in this simulation. One of the factors is completely lateralized (top left, highlighted in green), meaning that it loads only on ROIs in the left hemisphere. An additional illustration of this left-hemisphere factor is shown in Figure 5. The remaining 3 factors have both left- and right-hemisphere indicators.
2. The structure component matrix is a matrix with all 0 elements except on the secondary diagonal, i.e., the diagonal elements of the bottom left and top right quadrant are nonzero. The values of these secondary diagonals determine the strength of the symmetry.
3. The residual variance component matrix is a diagonal matrix where the elements are chosen such that the diagonal of **Σ**_*true*_ is **1**.

**Figure 5:**
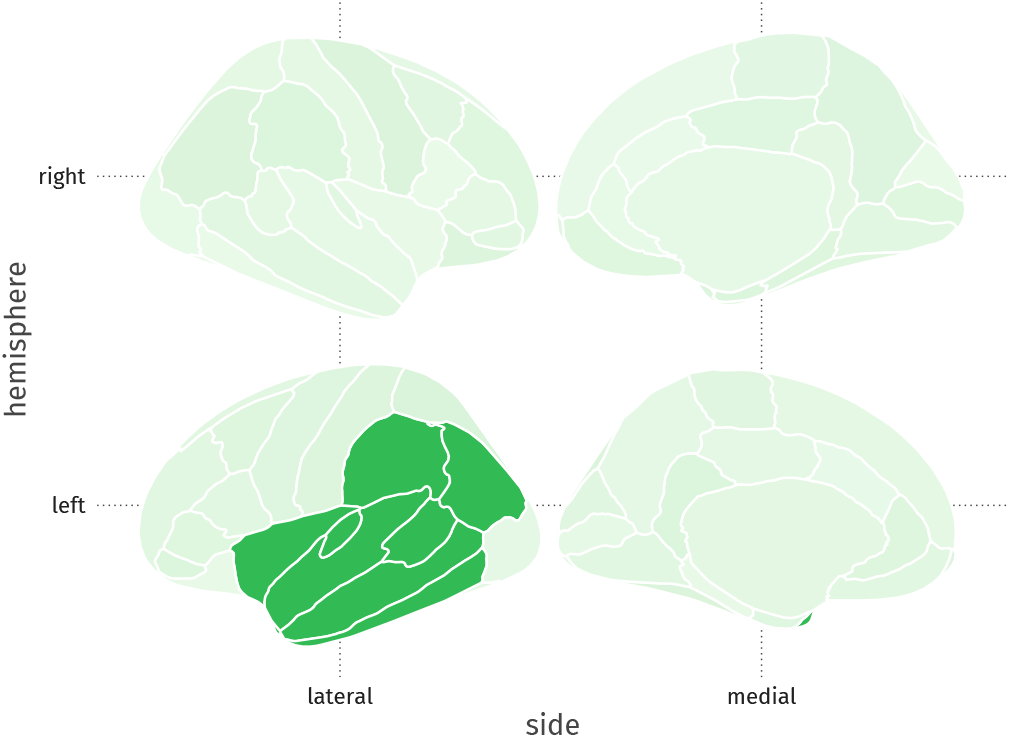
Example lateralized factor (the first factor in the simulation). Grey matter volume in 8 left-hemisphere regions of interest are predicted by the value on this factor.

For Sections 3.2 and 3.3, data were generated with a sample size of 650, a latent correlation of either 0 or 0.5, bilateral factor loadings of 0.5 or 0.7, lateral factor loadings of .425 or .595, and contralateral homology correlations of either 0 (pure EFA), 0.2 (minor symmetry), or 0.4 (major symmetry). These conditions were chosen to be plausible scenarios, similar to the observed data from Section 4. In each condition, 120 datasets were generated on which EFA and EFAST models with 4 factors were estimated. Thus, in each analysis the true number of factors is correctly specified before estimation. Section 3.4 explores different criteria for the choice of number of factors in the case of contralateral symmetry.

### 3.2 Effect of structured residuals on factor loadings

In this section, we compare estimated factor loadings from EFA and EFAST to the true factor loadings from the simulation’s data generating process. For each condition, 120 datasets were generated, to which both EFA and EFAST models were fit. The factor loading matrix for each model was then extracted, the columns reordered to best fit the true matrix, and the mean absolute error of the factor loadings per factor was calculated.

The results of these analyses are shown in 6. As hypothesized, allowing structured residuals affects how well the factor loadings are estimated from the datasets. Notably, as shown in 6 when performing regular EFA, the estimation error of the factor loadings increases when the symmetry becomes stronger, whereas the factor loading estimation error for the EFAST model remains at the level of regular EFA when there is no symmetry. Moreover, this effect is more pronounced in the lateralized factor as EFA tries to capture the symmetry within its factors: the lateralized factor becomes bilateral, leading to a larger error relative to the factors which are already bilateral, and an incorrect inference regarding the nature of the thus estimated factor. Although Figure 6 shows only the condition with factor loadings of 0.5 and factor covariance of 0.5, the pattern is similar for different factor loading strengths and with no factor covariance (see Appendix B).

**Figure 6:**
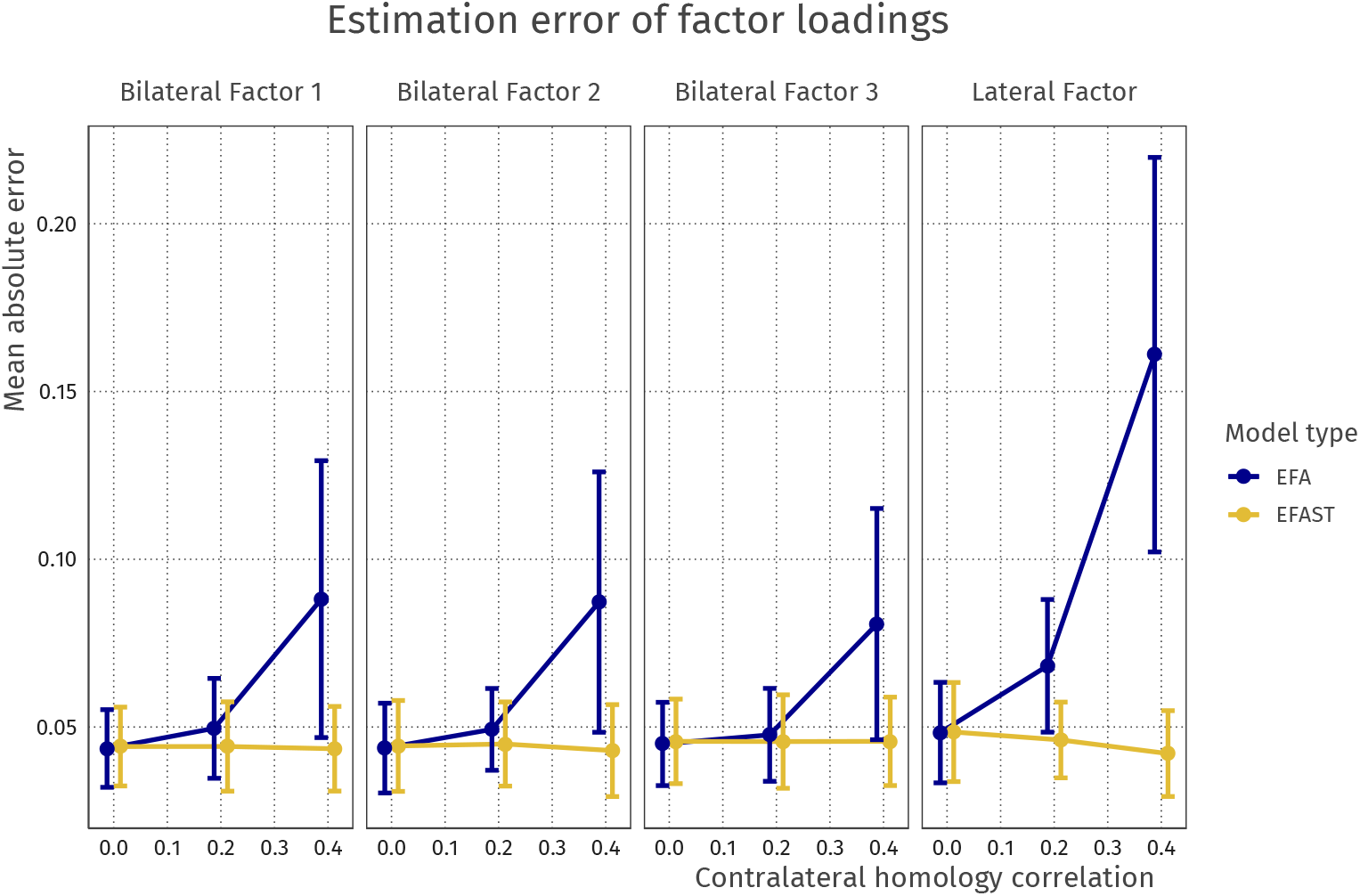
Mean absolute error for factor loadings of EFA versus EFAST models with increasing amounts of contralateral symmetry correlation. This plot comes from the condition where the covariance of the latent variables is 0.5, and the factor loadings are 0.5. The plot shows that for both bilateral and lateralised factors, EFA starts to exhibit more error as symmetry increases, more so for the lateral factor, whereas EFAST performance is nominal over these conditions. Error bars indicate 95% Wald-type confidence intervals.

Results from this section suggest that for the purpose of factor loading estimation, EFA and EFAST perform equally well in the case where a model without residual structure is the true underlying model, but EFAST outperforms EFA when residual structure in the observed data becomes stronger. In other words, implementing EFAST in the absence of residual structure does not seem to have negative consequences for estimation error, suggesting it may also be a useful default if a specific residual structure is thought, but not known, to exist. This is in line with Cole et al. (2007), who argue that in many situations including correlated residuals does not have adverse effects, but omitting them does.

### 3.3 Effect of structured residuals on factor covariances

Here, we compare how well EFA and EFAST retrieve the true factor covariance values. For both methods, we used geomin rotation with an epsilon value of 0.01 as implemented in `lavaan` 0.6.4 (Rosseel, 2019). The matrix product of the obtained rotation matrix ****H**** then represents the estimated factor covariance structure of the EFA factors: **Ψ**_*EFA*_ = ***H***^*T*^ ***H*** (Asparouhov and Muthén, 2009, eq. 22).

The mean of the off-diagonal elements of the **Ψ**_*EFA*_ matrix were then compared to the true value of 0.5 for increasing symmetry strength. The results are shown graphically in Figure 7. Here, it can be seen that with this rotation method the latent covariance is underestimated in all cases, although less so with stronger factor loadings. Furthermore, EFA performs worse as the symmetry increases, whereas the performance of EFAST remains stable regardless of the degree of contralateral homology, again suggesting no adverse effects to implementing EFAST in the absence of contralateral correlations. In the case of uncorrelated factors (not shown), the two methods perform similarly well.

**Figure 7:**
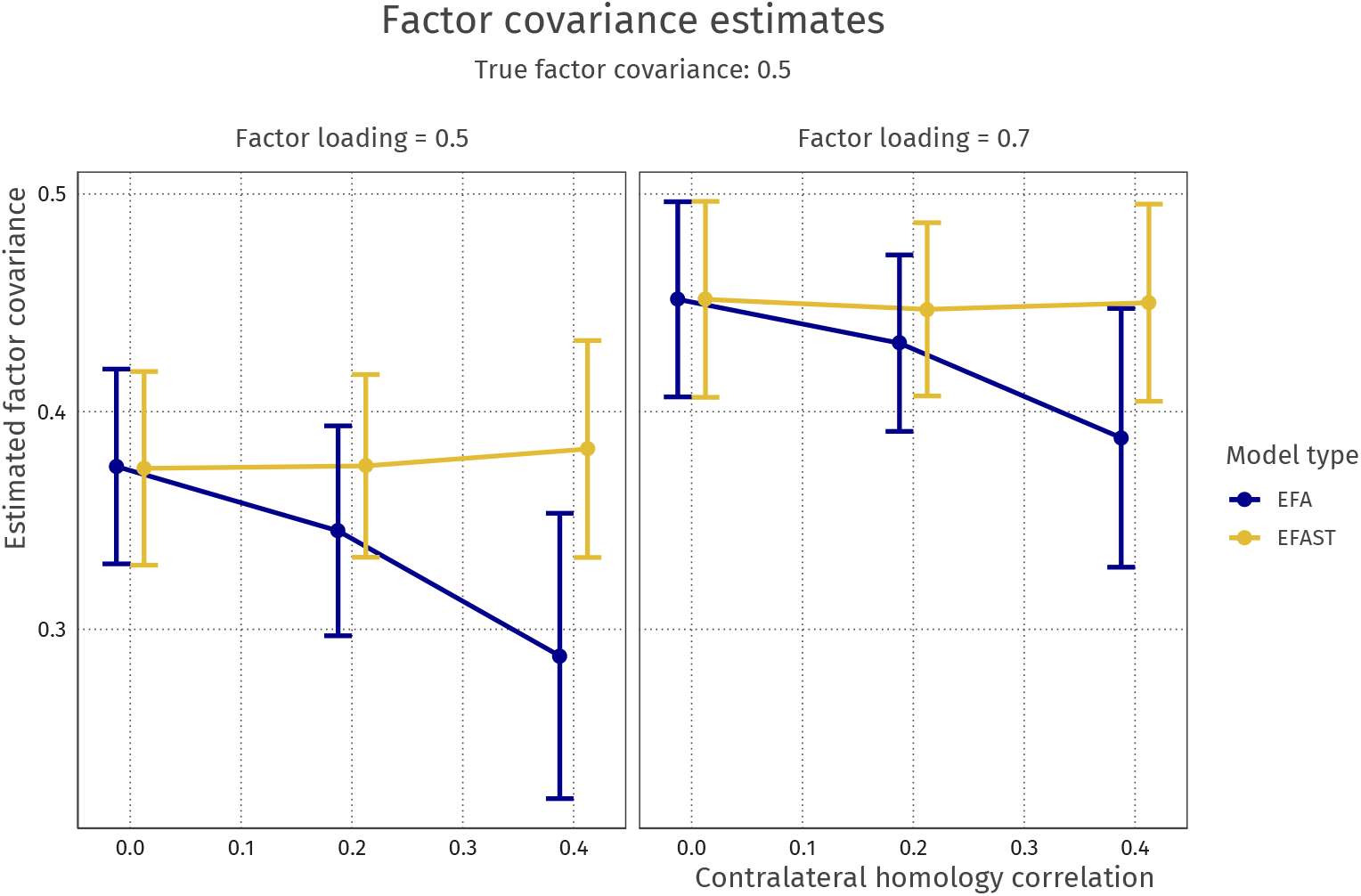
Latent covariance estimates for different levels of contralateral homology correlation. The true underlying latent covariance is 0.5; both methods underestimate the latent covariance but EFA becomes more biased as symmetry increases. Error bars indicate 95% Wald-type confidence intervals.

The results from this section shows that in addition to better factor recovery for EFAST, the recovery of factor covariance is also improved relative to EFA. Again, even when the data-generating mechanism does not contain symmetry, EFAST performs at least at the level of the EFA model. Note that the overall model fit in terms of AIC or BIC in this situation is better for the EFA model, as it has fewer parameters.

### 3.4 Effect of structured residuals on model fit

In the above analyses, the number of factors was specified correctly for each model estimation (using either EFA or EFAST). However, in empirical applications the number of factors will rarely be known beforehand, so has to be decided on the basis of some criterion. A common approach to extracting the number of factors, aside from computationally expensive strategies such as parallel analysis (Horn, 1965), is model comparison through information criteria such as the AIC or BIC (e.g. (Vrieze, 2012). In this procedure, models with increasing numbers of factors are estimated, and the best fitting model in terms of these criteria is chosen.

In this simulation, we generated 100 datasets as in Figure 4 – i.e., strong loadings and medium symmetry – and we fit EFA and EFAST models with 2 to 10 factors. Across these solutions we then compute the information criteria of interest. Here we choose the two most common information criteria (the AIC and BIC) as well as the sample-size adjusted BIC (SSABIC), as this is the default in the ESEM function of the `psych` package (Revelle, 2018). The results of this procedure are shown in Figure 8. Each point indicates a fitted model. The means of the information criteria are indicated by the solid lines.

**Figure 8:**
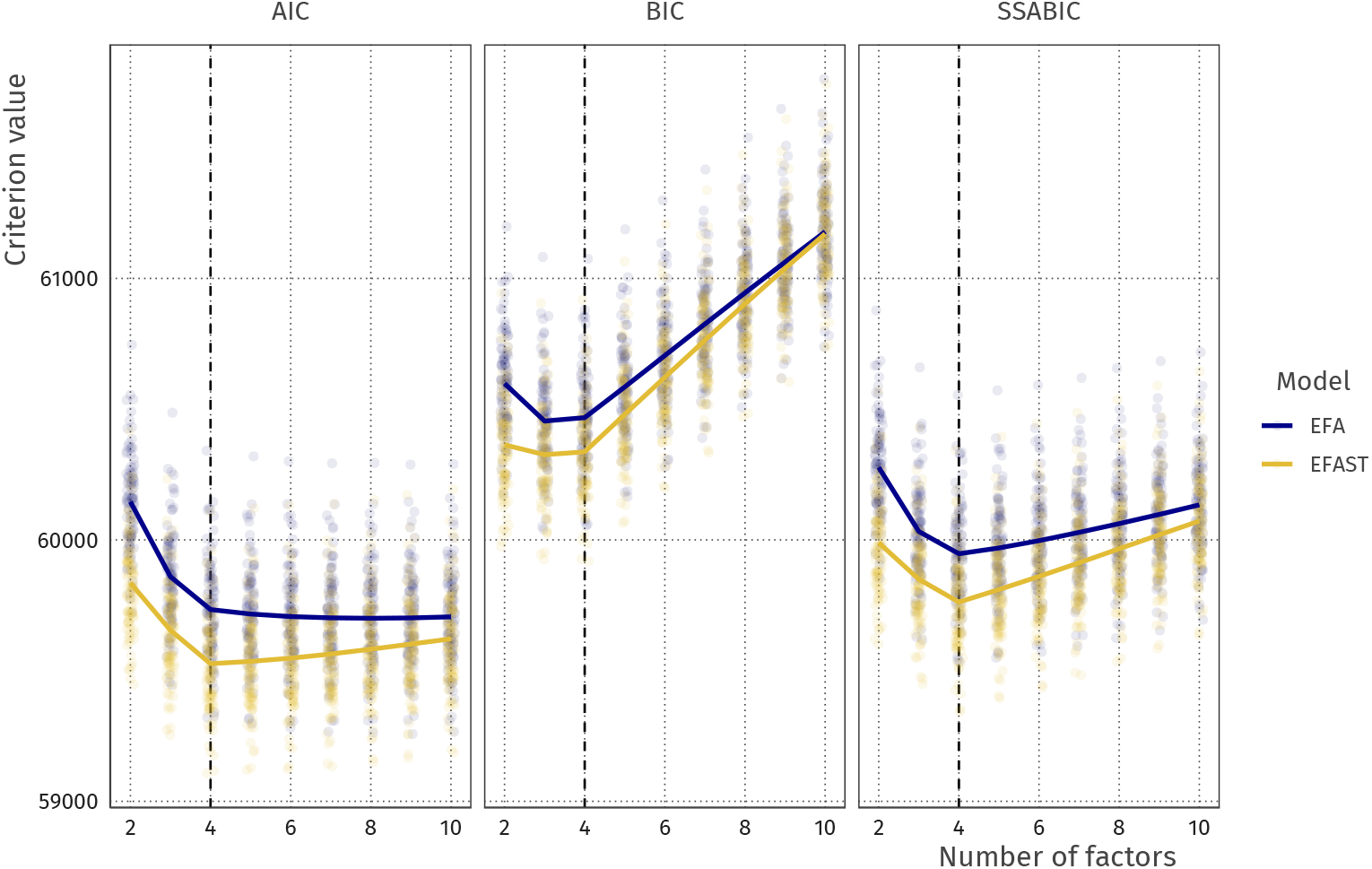
AIC and BIC values for increasing number of factors with EFA and EFAST models. Lines indicate expectations: the vertices are at the mean values for these criteria. The true number of factors is 4 (dashed vertical line).

The plot in Figure 8 shows that across all factor solutions, EFAST shows better fit than EFA, suggesting the improvement in model fit outweighs the additionally estimated parameters. In addition, the AIC tends to overextract factors, the BIC slightly underextracts, and the SSABIC shows the best extraction performance (see also Appendix C). In practice, therefore, we suggest using SSABIC when determining the number of factors and model fit is of primary concern. Note that a researcher may also wish to determine the number of factors based on other considerations, such as usability in further analysis, estimation tractability, or theory.

## 4 EFAST in practice: Modeling brain imaging data

In the field of cognitive neuroscience, a large body of work has demonstrated close ties between individual differences in brain structure and concurrent individual differences in cognitive performance such as intelligence tasks (e.g. van Basten et al., 2015). Moreover, different aspects of brain structure can be sensitive to clinical and pre-clinical conditions such as grey matter for multiple sclerosis (Eshaghi et al., 2018), white matter hyperintensities for cardiovascular factors (Fuhrmann et al., 2019) and white matter microstructure for conditions such as ALS (Bede et al., 2015), Huntingtons (Rosas et al., 2010) and many other conditions.

However, one perennial challenge in imaging is how to deal with the dimensionality of imaging data. Depending on the spatial resolution, a brain image can be divided into as many as as 100,000 individual regions, or voxels, rendering mass univariate approaches vulnerable to issues of multiple comparison. An alternative approach is to focus on anatomically defined sections called regions of interest or ROI’s. However, this only solves the challenge of dimensionality in part, by grouping adjacent voxels into meaningful regions. An emerging approach is therefore to take a covariance approach to such regions, by studying how neural measures in different regions covary across populations. This offers a promising strategy to reduce the high dimensional differences in brain structure into a tractable number of components, or factors, not limited by spatial adjacency. However, standard techniques such as EFA or PCA do not easily allow for the integration of a fundamental biological fact: That there exists strong contralateral symmetry between brain regions, such that any given region (e.g. the left lingual gyrus) is generally most similar to the same region on the other side of the brain. Here, we show how we can combine the strengths of exploratory data reduction with the integration of a priori knowledge about the brain into a more sensible, anatomically plausible factor structure which can either be pursued as an object of intrinsic interest or used as the basis for further investigations (e.g. which brain factors are most strongly associated with phenotypic outcomes).

### 4.1 Empirical example: Grey matter volume

#### 4.1.1 Data description

The data we use is drawn from the Cambridge Centre for Ageing and Neuroscience (Taylor et al., 2017; Shafto et al., 2014). Cam-CAN is a community derived lifespan sample (ages 18-88) of healthy individuals. Notably, the raw data from the Cam-CAN cohort is freely available through our data portal https://camcan-archive.mrc-cbu.cam.ac.uk/dataaccess/. The sample we discuss here is based on 647 individuals. For the purposes of this project we use morphometric brain measures derived from the T1 scans. Specifically, we used the Mindboggle pipeline (Klein et al., 2017) to estimate region based grey matter volume, using the underlying freesurfer processing pipeline. To delineate the regions, we here use the Desikan-Killiany-Tourville atlas for determining the ROIs (Klein and Tourville, 2012) as illustrated in Figure 9.

**Figure 9:**
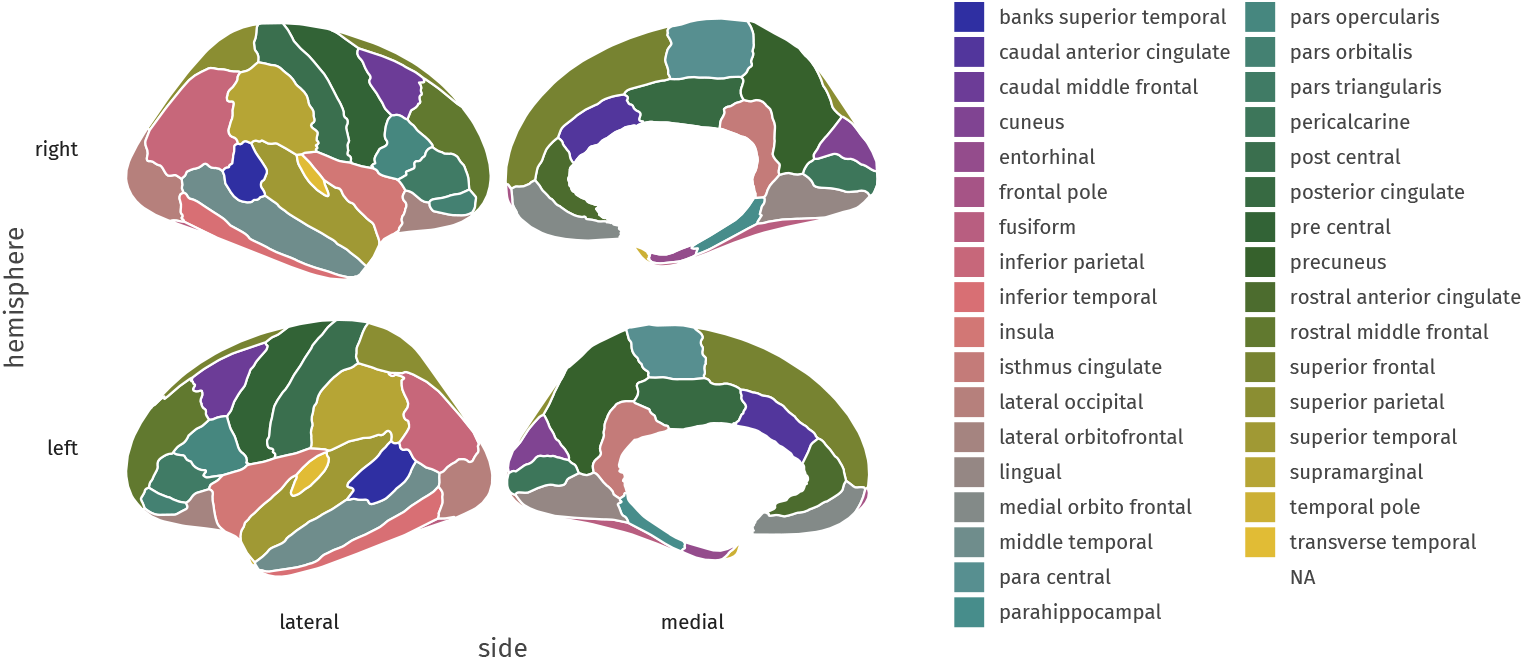
Desikan–Killiany–Tourville atlas used in the empirical illustration, as included in the `ggseg` package (Mowinckel and Vidal-Piñeiro, 2019).

We focus only on grey matter (not white matter) and only on cortical regions (not subcortical or miscellaneous regions such as ventricles) with the above atlas, for a total of 68 brain regions. The correlation matrix of regional volume metrics is shown in Figure 10, where the first 34 variables are regions of interest (ROIs) in the left hemisphere, and the last 34 variables are ROIs in the right hemisphere. The presence of higher covariance due to contralateral homology is clearly visible in the darker secondary diagonal ‘stripes’ which show the higher covariance between the left/right version of each anatomical region. This is because the ROIs have the same order in both hemispheres, meaning that variable 1 and 34 are homologues, and 2 and 35, and so forth. Our goal is to reduce this high-dimensional matrix into a tractable set of ‘brain factors’, which we may then use in further analyses, such as differences in age sensitivity, in a way that respects known anatomical constraints.

**Figure 10:**
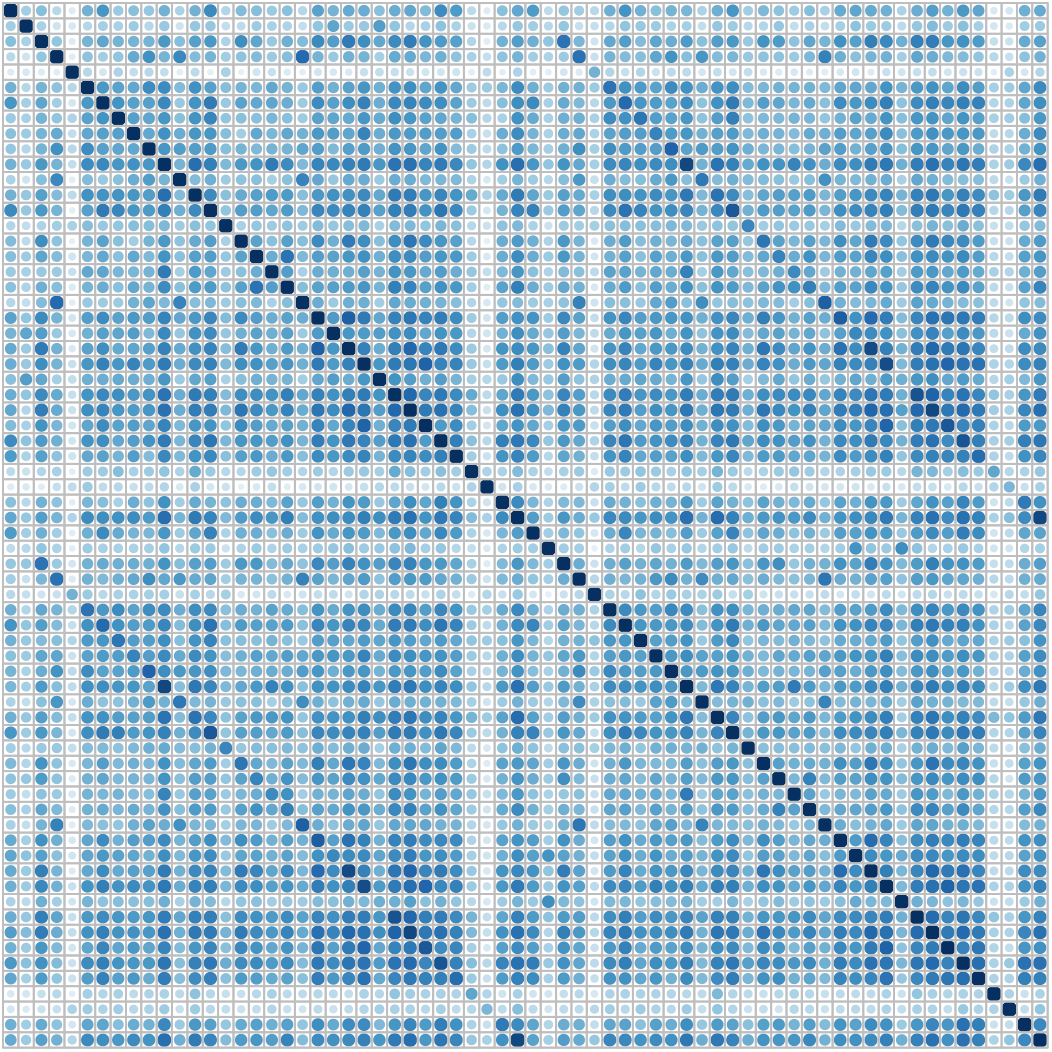
Correlation plot of cortical grey matter volume in 647 T1 weighted images of the Cam-CAN sample, estimated through Mindboggle in 34 brain regions in each hemisphere according to DKT segmentation. Darker blue indicates stronger positive correlation. Secondary diagonal lines are visible indicating correlation due to contralateral homology.

The default estimation using EFA will attempt to account for the strong covariance among homologous regions seen in this data, meaning it is unlikely for, say, the left insula and the right insula to load on different factors, and/or for a factor to be characterized only/mostly by regions in one hemisphere. To illustrate this phenomenon, we first run a six-factor, geomin-rotated EFA for the above data. The factor loadings for each ROI in the left and right hemispheres are plotted in Figure 11. A strong factor loading for a ROI in the left hemisphere is likely to have a strong factor loading in the right hemisphere due to the homologous correlation, as shown by the strong correlations for each of the factors.

**Figure 11:**
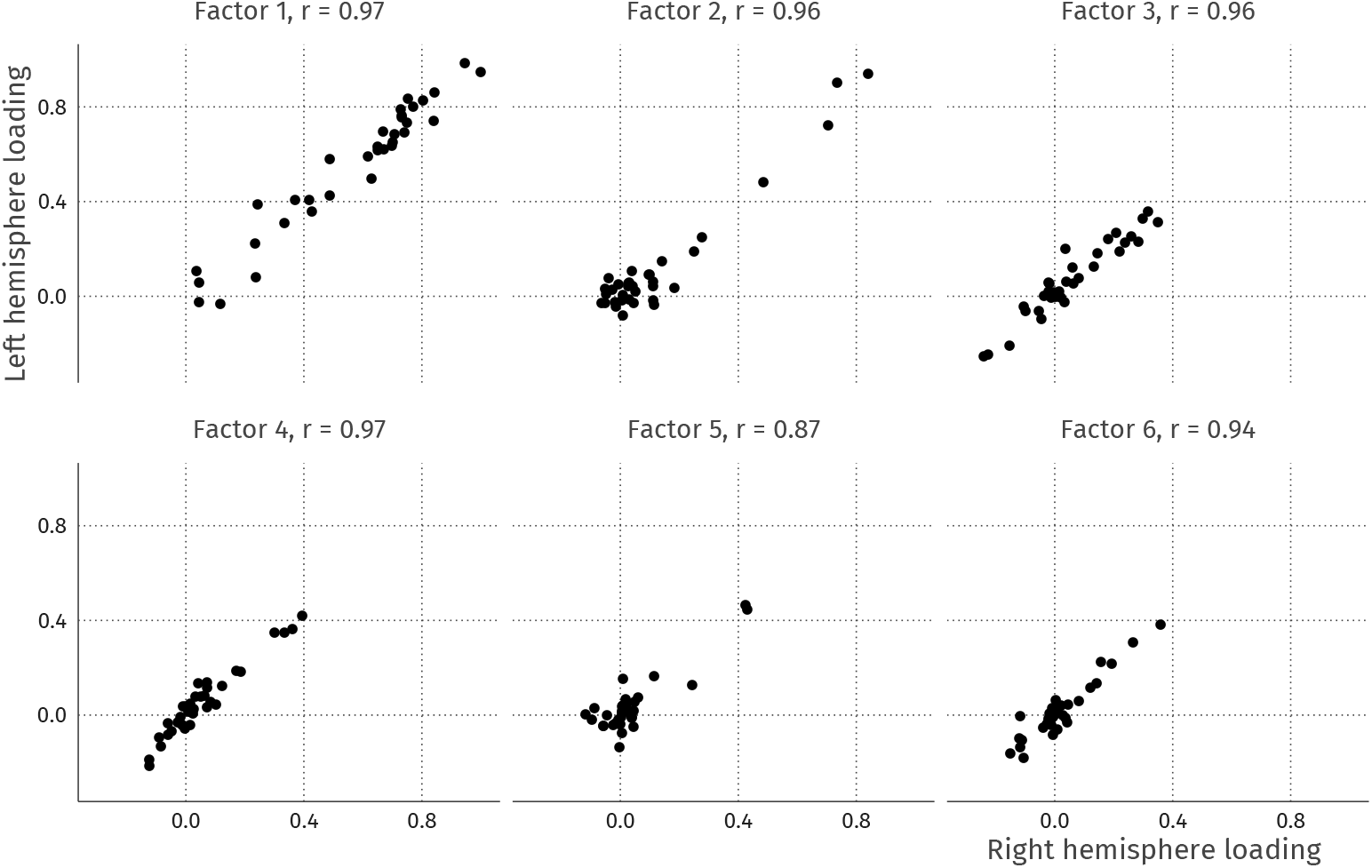
Left-right hemisphere factor loading correlations. The correlations between the loadings are high, indicating a strong similarity between the loadings in the left and right hemispheres.

In EFA, the resulting factors thus inevitably capture correlation due to contralateral symmetry, inflating or deflating factor loadings due to these contralateral residual correlations. Most problematically from a substantive neuroscientific standpoint, this distortion means it is effectively impossible to discover *lateralized factors*, i.e. patterns of covariance among regions expressed only, or dominantly, in one hemisphere. This is undesirable, as there is both suggestive and conclusive evidence that some neuroscientific mechanisms may display asymmetry. For instance, typical language ability is associated with an asymmetry in focal brain regions (e.g., Gauger et al., 1997; Bishop, 2013), whereas structural differences in the right hemisphere may be more strongly associated with face perception mechanisms (Frässle et al., 2016). Developmentally, there is evidence that the degree of asymmetry changes may lead to changes in asymmetry (e.g. (Plessen et al., 2014). Within a SEM context, recent work shows that model fit of a hypothesized covariance structure may differ substantially between the right and left hemispheres despite when focusing on the same brain regions (Meyer et al., 2019). The ignorance of traditional techniques for the residual structure may cause lateralized covariance factors to appear symmetrical instead, or to not be observed at all.

#### 4.1.2 Results

In this section, we compare the model fit and factor solutions of EFA and EFAST for the Cam-CAN data, and we show how EFAST decomposes the correlation matrix in Figure 10 into factor, structure, and residual variance components. The full annotated analysis script to reproduce these results is available as supplementary material to this manuscript.

The model estimation, shown in Figure 12, demonstrates that using common information criteria, the EFAST model performs considerably better than standard EFA. The BIC criterion, combined with the convergence of the models to an admissible solution, suggests that 6 factors is optimal for this dataset. While both AIC and SSABIC show that more factors may be needed to properly represent the data, we see that this quickly leads to nonconvergence. We here consider 6 factors to be a tractable number for further analysis. First and foremost, this 6-factor solution shows a much better model solution under EFAST (BIC ≈ 87500) than under EFA (BIC ≈ 90000), emphasizing the empirical benefits of appropriately modeling known biological constraints. Additionally, statistical model comparison through a likelihood ratio test shows that the EFAST model fits significantly better (see Table 1). Other fit measures such as CFI, RMSEA, and SRMR paint a similar story. The full factor loading matrix for both EFAST and EFA are shown in Appendix C.

**Figure 12:**
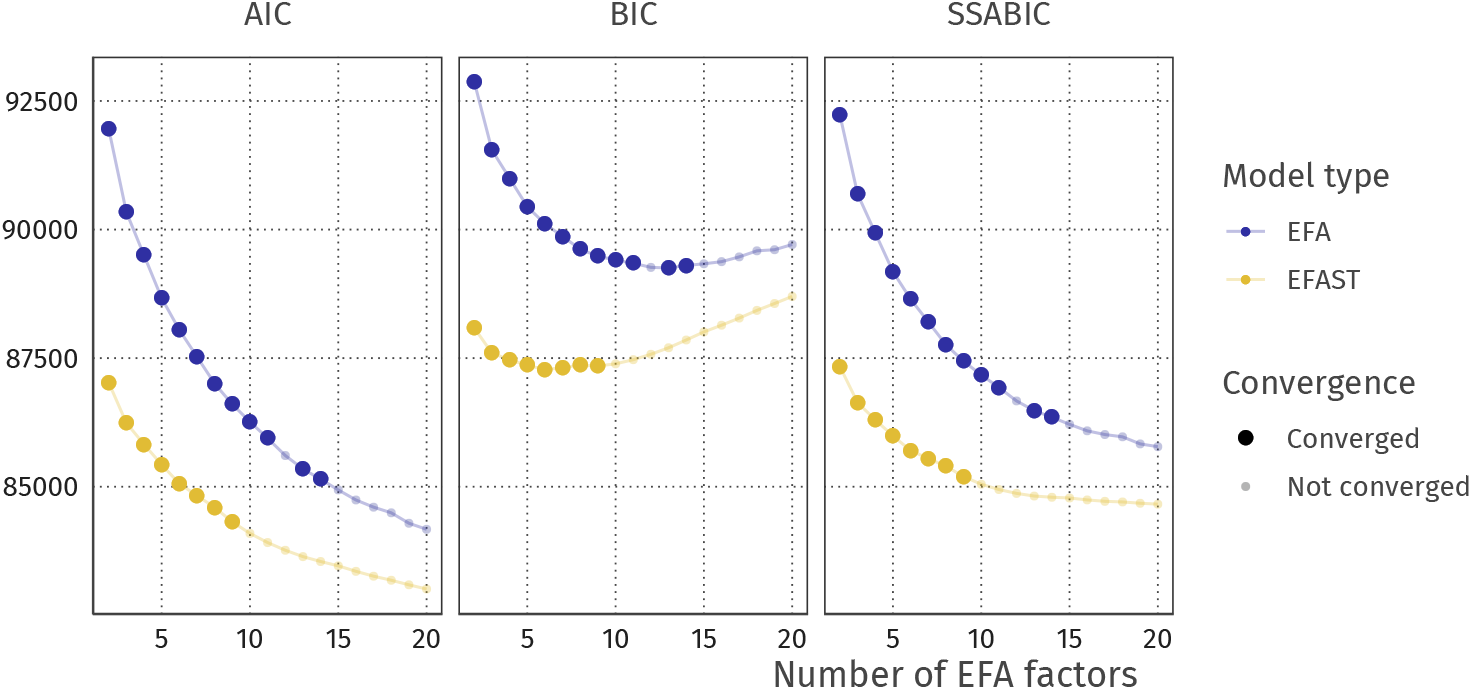
AIC and BIC for the with increasing numbers of EFA factors. Semitransparent points indicate models which are inadmissible either due to nonconvergence or convergence to a solution with problems (e.g., Heywood cases). In these cases we plot the information criteria based on the log-likehood computed at the time the estimation terminated.

**Table 1:** Comparing the fit of the EFAST and EFA models with 6 factors, using a likelihood ratio test and several fit criteria.

| | CFI | RMSEA | SRMR | $\chi^2$ | Df | $\Delta\chi^2$ | $\Delta Df$ | $\Pr(> \chi^2)$ |
| --- | --- | --- | --- | --- | --- | --- | --- | --- |
| EFAST | 0.912 | 0.057 | 0.209 | 5762.676 | 1851 |  |  |  |
| EFA | 0.843 | 0.075 | 0.342 | 8818.146 | 1885 | 3055.471 | 34 | < .001 |

The EFAST model decomposes the observed correlation matrix from Figure 10 into the three components displayed in Figure 13. The most notable observation here is the separation of symmetry structure (last panel) and latent factor-implied structure (first panel): the factor solution (first panel) does not attempt to explain the symmetry structure seen in the data (i.e. the characteristic diagonal streaks are no longer present). This indicates that the EFAST model correctly separates symmetry covariance from underlying trait covariance in real-world data.

**Figure 13:**
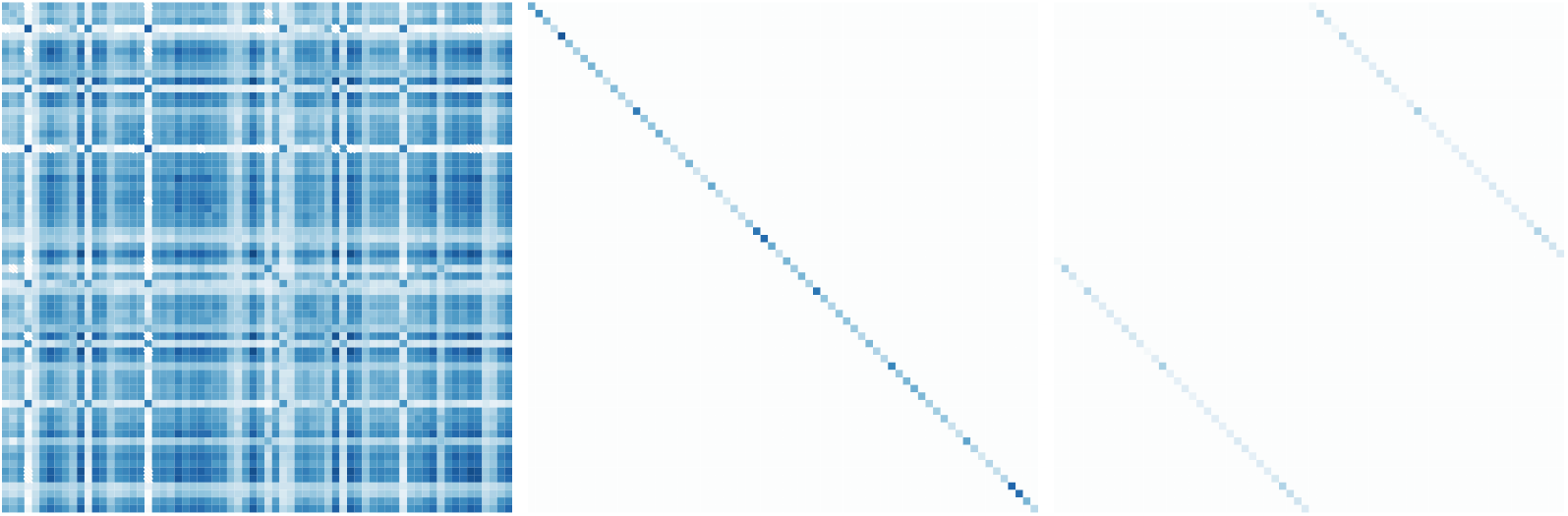
Extracted correlation matrix components using a 6-factor EFAST model with unconstrained correlations. From left to right: factor-implied correlations, residual variance, and structure matrix.

We also extracted the estimated factor covariance, shown as a network plot in Figure 14. For EFA, some latent variables show very strong covariance, clustering them together due to the contralateral symmetry. This effect is not visible in the EFAST model, which shows a more well-separated latent covariance structure. This suggests that one consequence of a poorly specified EFA can be the considerable overestimation of factor covariance, which in turn adversely affects the opportunities to understand distinct causes or consequences of individual differences in these factors.

**Figure 14:**
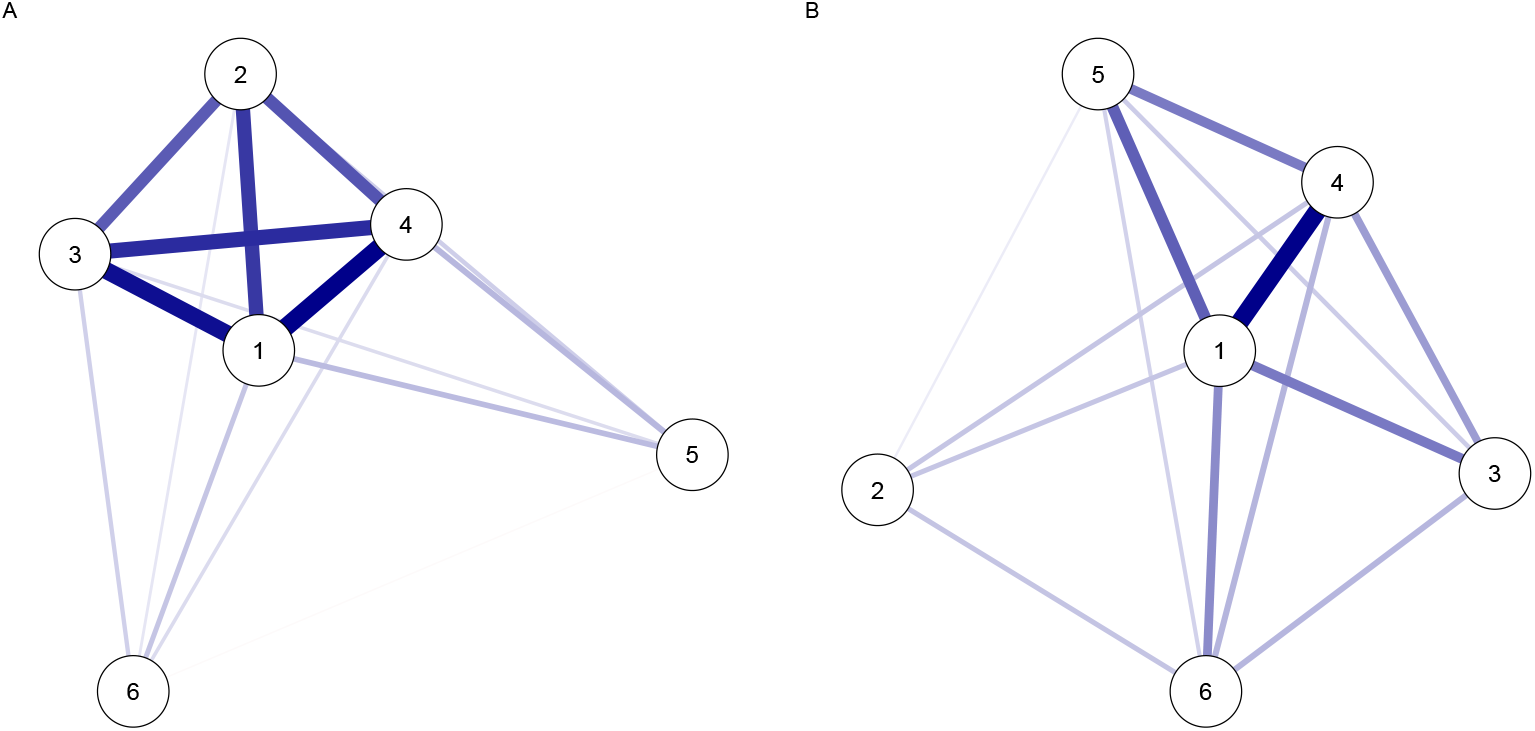
Network plots of the latent covariance for EFA (panel A) and EFAST (panel B).

**Figure 15:**
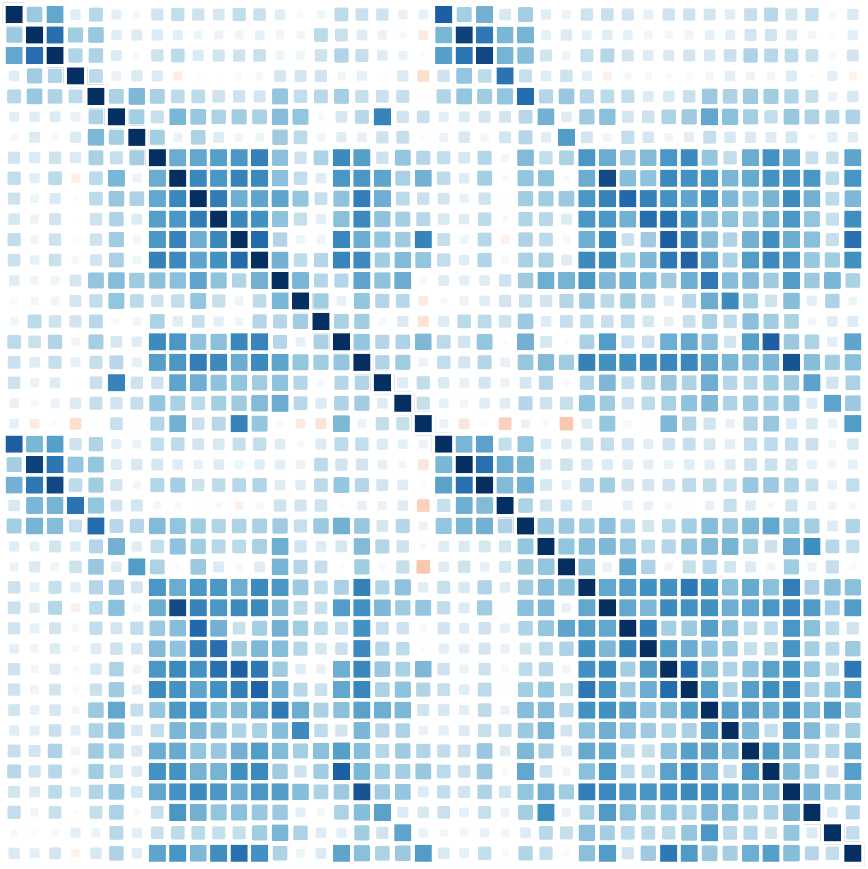
Correlation matrix for Cam-CAN white matter tractography data (fractional anisotropy).

### 4.2 Empirical example: White matter microstructure

#### 4.2.1 Data description

Our second empirical example uses white matter structural covariance networks. We use 42 tracts from the ICBM-DTI-81 atlas (Mori et al., 2008), including only those tracts with atlas-separated left/right tracts (i.e. excluding divisions of the corpus callosum – For a full list, see appendix). As anatomical metric we use tract-based mean fractional anisotropy, a summary metric sensitive (but not specific) to several microstructural properties ((Jones et al., 2013). For more details regarding the analysis pipeline, see (Kievit et al., 2016). The same tracts and data were previously analysed in (Jacobucci et al., 2019).

#### 4.2.2 Results

We chose 6 factors for the EFAST and EFA models based on the SSABIC in combination with the convergence limitations. In Table 2, the two models are compared on various characteristics. From the likelihood ratio test, we can see that the EFAST model represents the white matter data significantly better (*χ*^2^(21) = 3632.586, *p* < .001), and inspecting the SSABIC values (EFA = 59120, EFAST = 55727) leads to the same conclusion. In addition, the CFI, RMSEA, indicate better fit for the EFAST model, too.

**Table 2:**
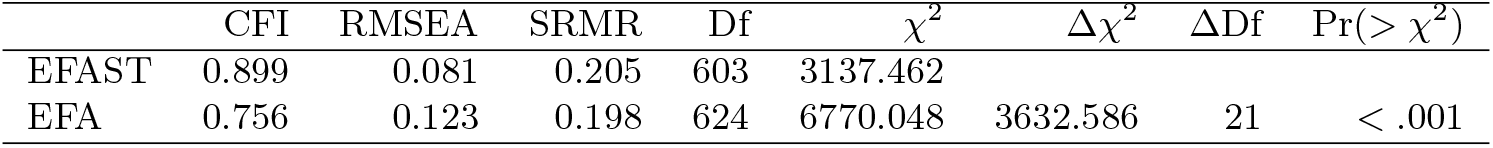
Comparing the fit of the EFAST and EFA models with 6 factors for the white matter data, using a likelihood ratio test and several fit criteria.

| | CFI | RMSEA | SRMR | Df | $\chi^2$ | $\Delta\chi^2$ | $\Delta Df$ | $\Pr(> \chi^2)$ |
| --- | --- | --- | --- | --- | --- | --- | --- | --- |
| EFAST | 0.899 | 0.081 | 0.205 | 603 | 3137.462 |  |  |  |
| EFA | 0.756 | 0.123 | 0.198 | 624 | 6770.048 | 3632.586 | 21 | < .001 |

The only index which indicates slightly poorer fit is the SRMR. The difference is very small in this case, but nonetheless it is relevant to show where these differences lie. A visual representation of the root square residual (observed - implied) correlations – which form the basis of the SRMR fit index – can be found in Figure 16. The figure shows that EFAST is able to represent the symmetry better: it has almost no residuals on the secondary diagonals. The remaining residuals are very similar, though slightly higher, leading to a higher SRMR.

**Figure 16:**
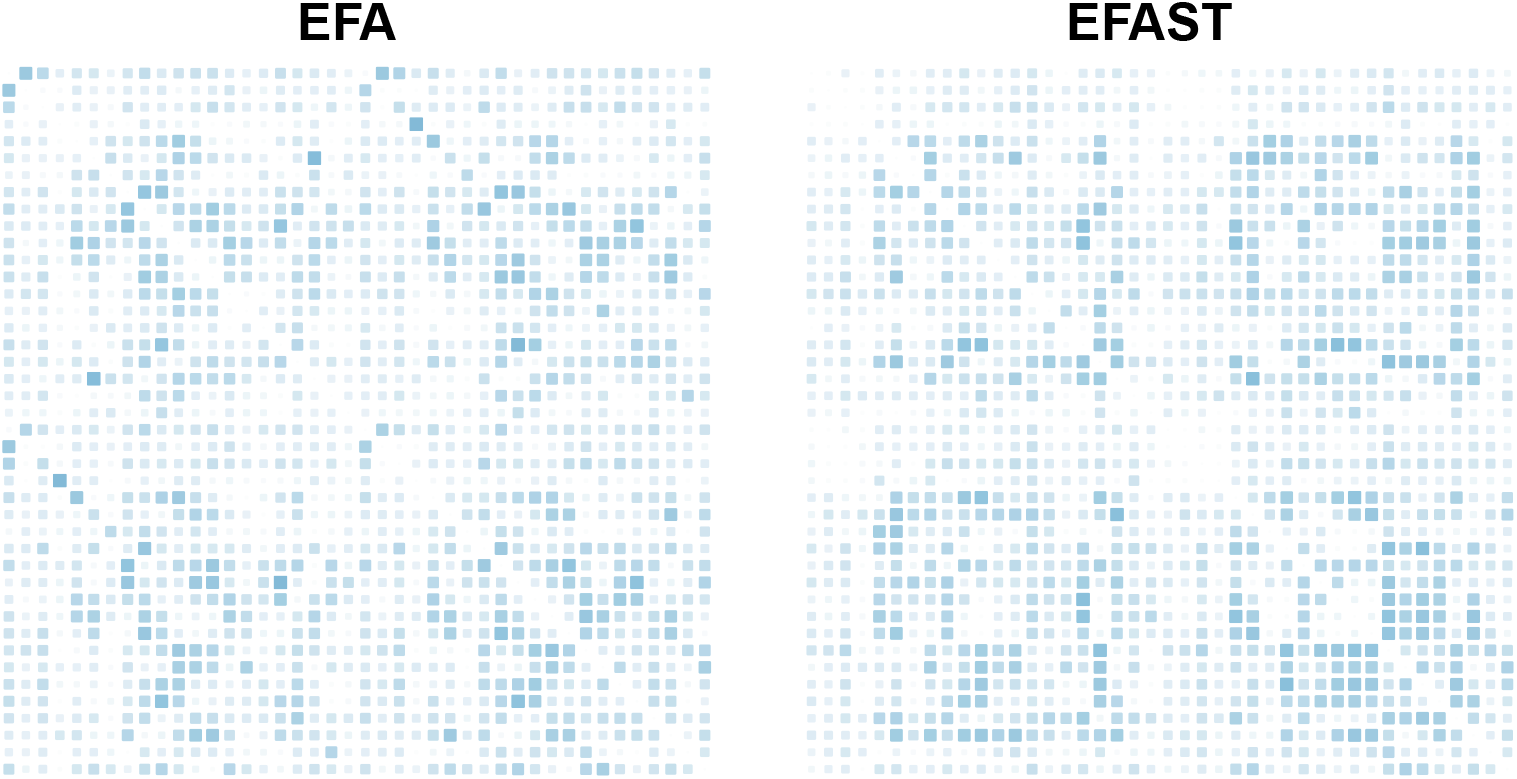
Visual representation of the root square residual (observed - implied) correlations, which form the basis of the SRMR fit index.

### 4.3 Empirical example: Resting state Functional connectivity

#### 4.3.1 Data description

Our previous examples correlation matrices capturing between-individual similarities across regions. However, the same techniques can be implemented at the within-subject level given suitable data. One such measure is *functional connectivity* which reflects the temporal connectivity between regions during rest or a given task, and captures the purported strength of interactions, or communications, between regions (Van Den Heuvel and Pol, 2010). Here we use functional connectivity matrices from 5 participants in the Cam-CAN study measured during an eyes-closed resting state block. We focus on 90 cortical and sub-cortical regions from the AAL-atlas (Tzourio-Mazoyer et al., 2002). The methodology to compute the connectivity metrics is outlined in (Geerligs et al., 2017), and the data reported here have been used in (Lehmann et al., 2019).

#### 4.3.2 Results

For this example, data from the first participant was used to perform the model fit assessments. We performed a similar routine as with the previous empirical datasets for determining the number of factors: we fit the EFAST and EFA models for 2-16 factors and compare their information criteria. All of the models converged, and the optimal model based on the BIC is a 13-factor EFAST model.

The 13-factor EFAST model was then compared to the 13-factor EFA model on various fit indices. The results of this comparison can be found in Table 3. Across the board, the EFAST model has better fit, as the EFAST CFI, RMSEA, SRMR and *χ*^2^ fit indices outperform those for the EFA model, demonstrating that accounting for the bilateral symmetry in dimension reduction through factor analysis leads to better fitting model of the data.

**Table 3:** Comparing the fit of the EFAST and EFA models with 13 factors for the functional resting state data, using a likelihood ratio test and several fit criteria.

| | CFI | RMSEA | SRMR | Df | $\chi^2$ | $\Delta\chi^2$ | $\Delta Df$ | $\Pr(>\chi^2)$ |
| --- | --- | --- | --- | --- | --- | --- | --- | --- |
| EFAST | 0.836 | 0.093 | 0.253 | 2868.000 | 9350.278 |  |  |  |
| EFA | 0.774 | 0.108 | 0.272 | 2913.000 | 11828.126 | 2477.848 | 45.000 | 0.000 |

This approach also allows for comparing the factor loadings for the different participants. For illustration, the plot in Figure 18 shows the profile of factor loadings for the first three factors (columns) across the five participants (rows). These profile plots can be a starting point for comparison of the connectivity structure across participants, where higher correlation among participants means a more similar connectivity structure, while taking into account the symmetry in the brain. For example, for Factor 1, participant 3 has a quite different functional connectivity factor loading profile than the other participants.

**Figure 17:**
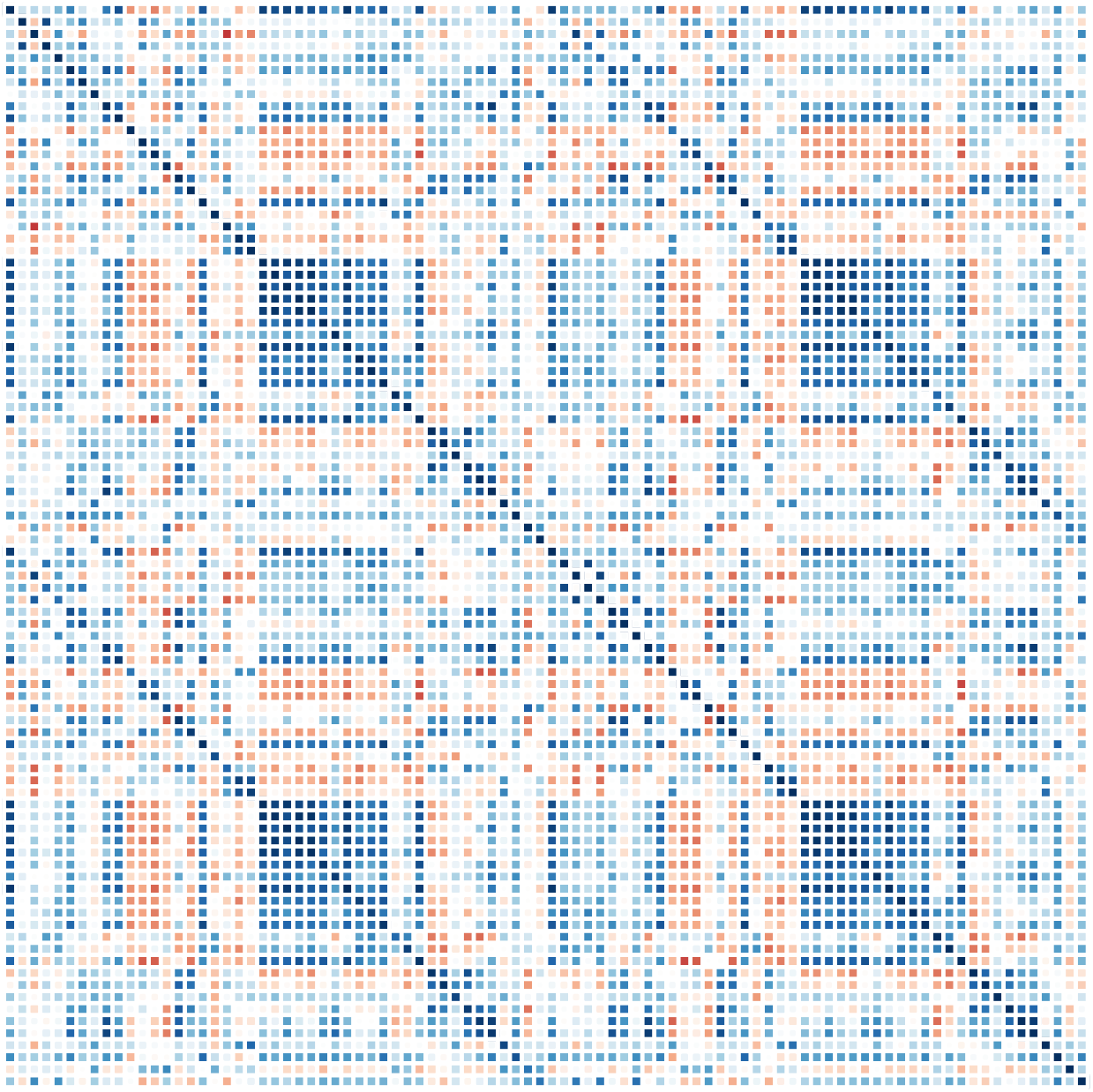
Correlation matrix for the first participant in the Cam-CAN resting state functional connectivity dataset.

**Figure 18:**
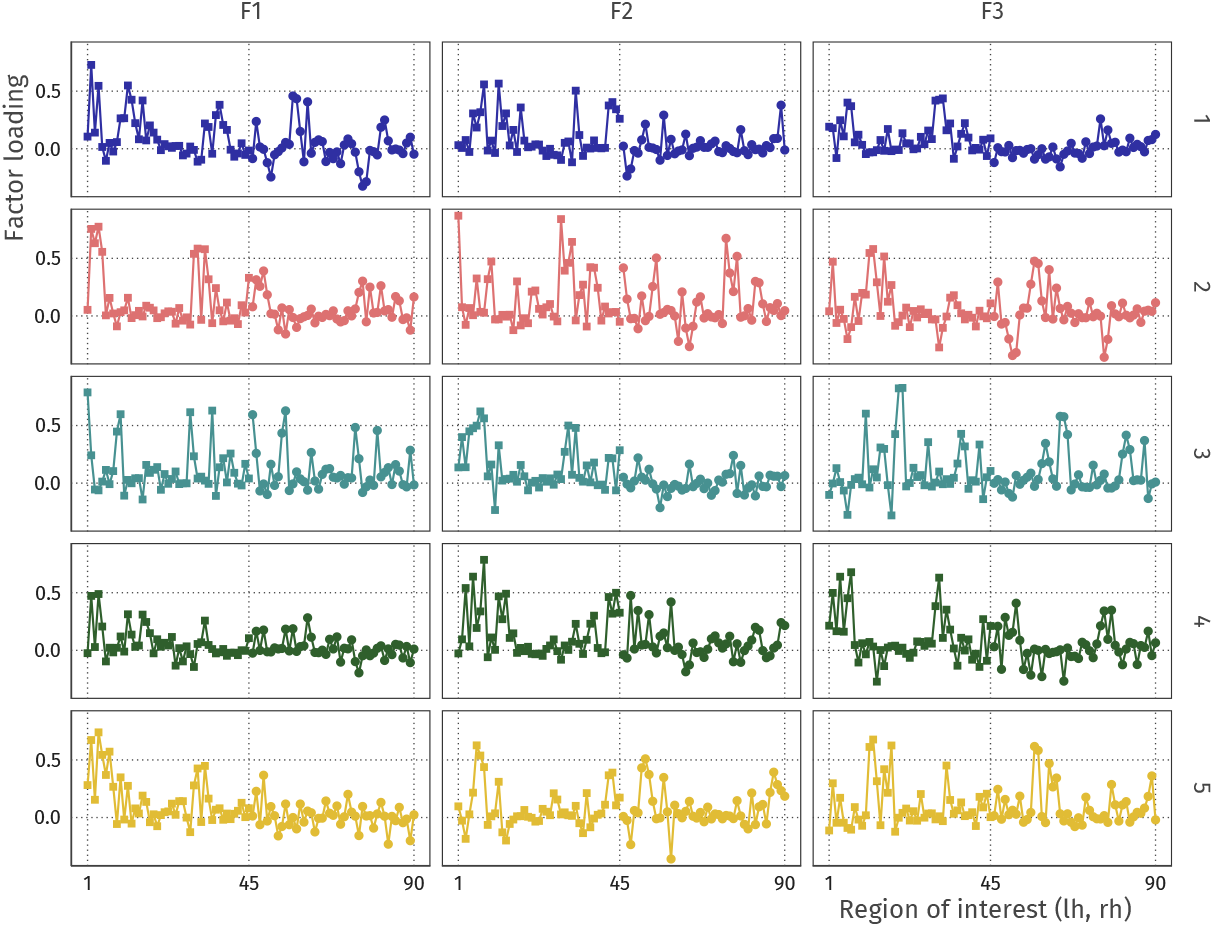
Comparison of factor loading profiles for the first three factors (columns) across five participants (rows).

## 5 Model-based lateralization index

In Section 3 we showed how the EFAST approach yields a more veridical representation of the factor structure than EFA. However, using EFAST yields an additional benefit: our model allows for estimating the extent of symmetry in each ROI, while taking into account the overall factor structure. This enables researchers to use this component of the analysis for further study. The (lack of) symmetry may be of intrinsic interest, such as in language development research (Schuler et al., 2018), intelligence in elderly (Moodie et al., 2019), and age-related changes in cortical thickness asymmetry (Zhou et al., 2013; Plessen et al., 2014). In the `efast` package, we have implemented a specific form of lateralization which is based on a variance decomposition in the ROIs. Our lateralization index (LI) represents the proportion of residual variance (given the trait factors) in an ROI that cannot be explained by symmetry. The index value is 0 if the bilateral ROIs are fully symmetric (conditional on the trait factors), and 1 if there is no symmetry.

The LI for each ROI in the grey matter volume example is shown in Figure 19. Here, we can see that there is high lateralization in the superior temporal sulcus and medial orbitofrontal cortex, but high symmetry in the lateral orbitofrontal cortex and the insula. In Figure 20, we additionally show in the white matter example that LI can naturally be supplemented by standard errors and confidence intervals. Thus, the EFAST procedure not only improves the factor solution under plausible circumstances for such datasets, but in doing so yields an intrinsically interesting metric of symmetry.

**Figure 19:**
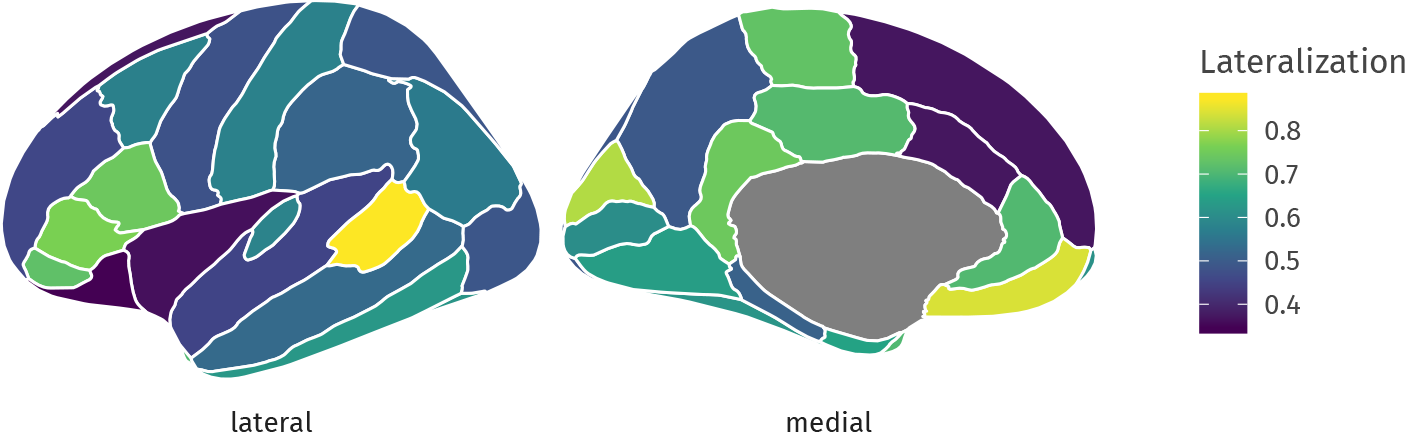
Amount of grey matter volume asymmetry per ROI. Dark blue areas are highly symmetric given the previously estimated 6-factor solution, and bright yellow areas are highly asymmetric. Such plots can be made and compared for different groups and statistically investigated for differences in symmetry for a common factor solution. A lateralization index (LI) of 0.8 means that 20% of the residual variance in grey matter volume in can be explained by symmetry.

**Figure 20:**
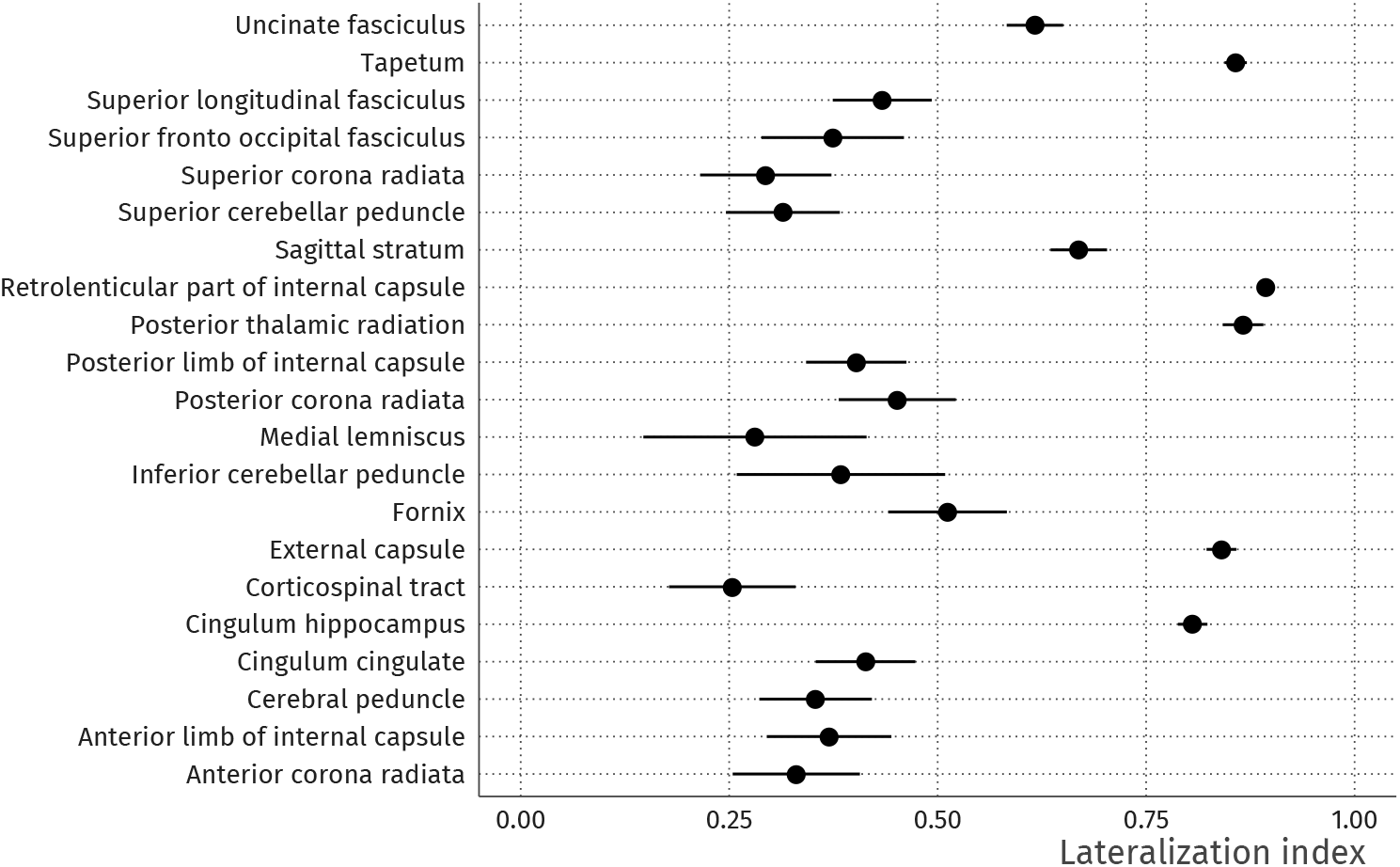
White matter lateralization index for a selected set of regions given the previously estimated 6-factor solution. Lower values means that bilateral ROIs are more symmetric conditional on the trait factors, higher values that they are less so. The line ranges indicate 95% confidence intervals, computed as *LI* ± 1.96 × *SE_LI_*, where the standard error *SE_LI_* is computed using the delta method.

## 6 Summary and discussion

In this paper, we have developed and implemented EFAST, a method for performing dimension reduction on data with residual structure. We show how this new method outperforms standard EFA across three separate datasets, by taking into account hemispheric symmetry. We have argued through both simulations and real-world data analysis that our method is an improvement in the dimension reduction step of such high-dimensional, structured data, yielding a more veridical factor solution. Such a factor solution can be the basis for further analysis, such as an extension of the factor model to prediction of continuous phenotype variables such as intelligence scores, or the comparison among different age groups. These extensions will be improved by building on a factor solution appropriately which takes into account the symmetry of the brain.

Care is needed in the interpretation of the factor solution as underlying dimensions, as the empirical application has shown that the absolute level of fit for both the EFA and EFAST models is not optimal. In addition, estimation of more complex factor models may lead to nonconvergence or inadmissible solutions. Such problems would need to be further investigated, potentially leading to more stable estimation, for example through a form of principal axis factoring, or potentially through penalization of SEM (Jacobucci et al., 2019; van Kesteren and Oberski, 2019). However, these limitations hold equally for EFA, and when comparing both methods it is clear from the results in this paper that the inclusion of structured residuals greatly improves the representation of the high-dimensional raw data by the low-dimensional factors. In summary, this relatively simple but versatile extension of classical EFA may be of considerable value to applied researchers with data that posses similar qualities to those outlined above. We hope our tool will allow those researchers to easily and flexibly specify and fit such models.

## Acknowledgements

We would like to thank Yves Rosseel for valuable input and the development of key tools, Linda Geerligs for providing the functional connectivity data, and Jonathan Helm and Øystein Sørensen for their helpful comments on an earlier version of this manuscript. E.-J. van Kesteren is supported by the Netherlands Organization for Scientific Research (NWO) under grant number 406.17.057. R. A. Kievit is supported by the and the UK Medical Research Council SUAG/047 G101400. This project received funding from the European Union’s Horizon 2020 research and innovation programme (grant agreement number 732592).

## A Symmetry pattern recovery with default EFA

**Figure 21:**
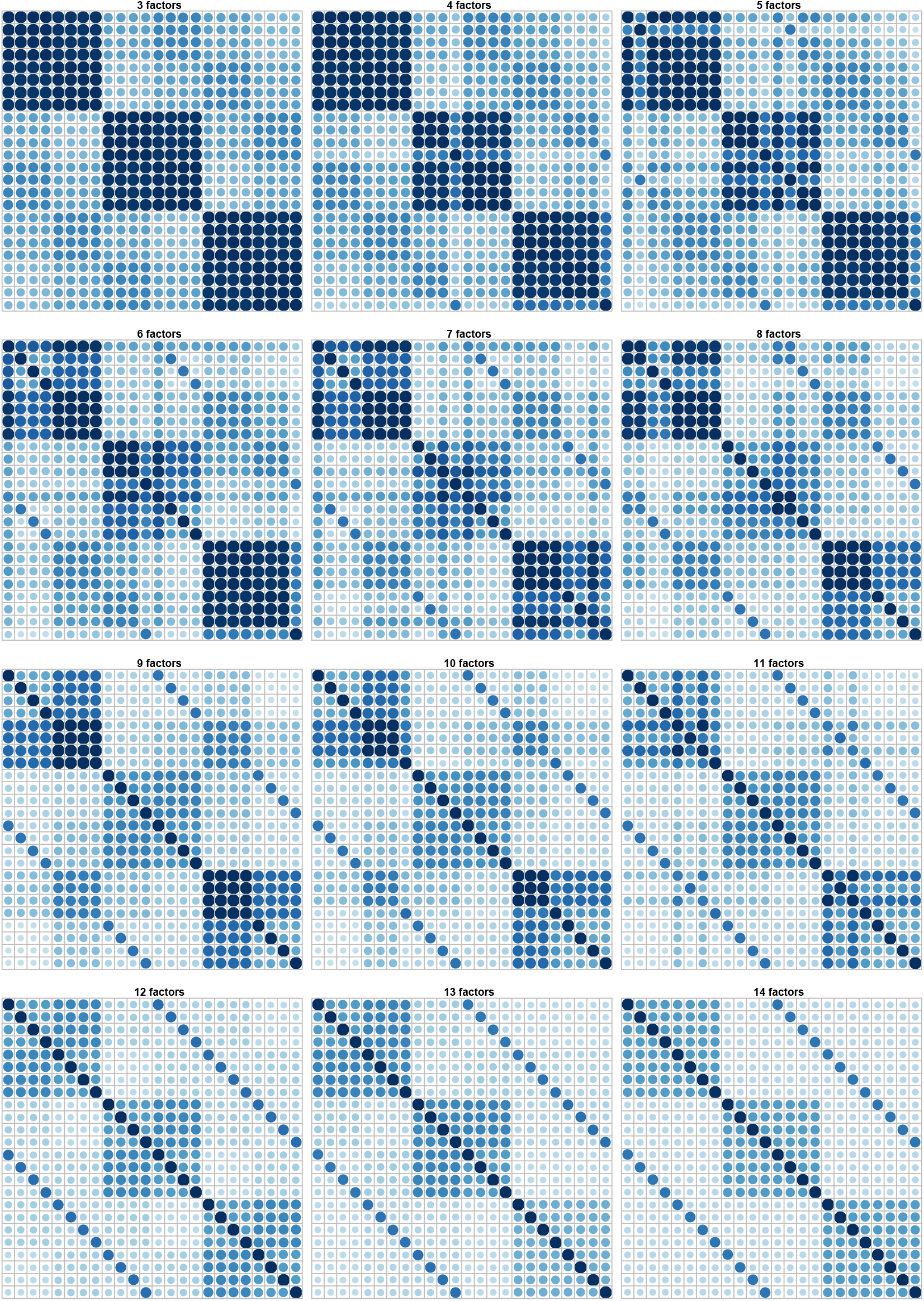
Predicted correlation matrix for EFA models with *M* factors for the example observed correlation matrix of Section 2, Figure 1. Proper recovery of the observed pattern happens around 12 factors (bottom left frame).

## B Comparing EFA and EFAST in factor loading estimation error

**Figure 22:**
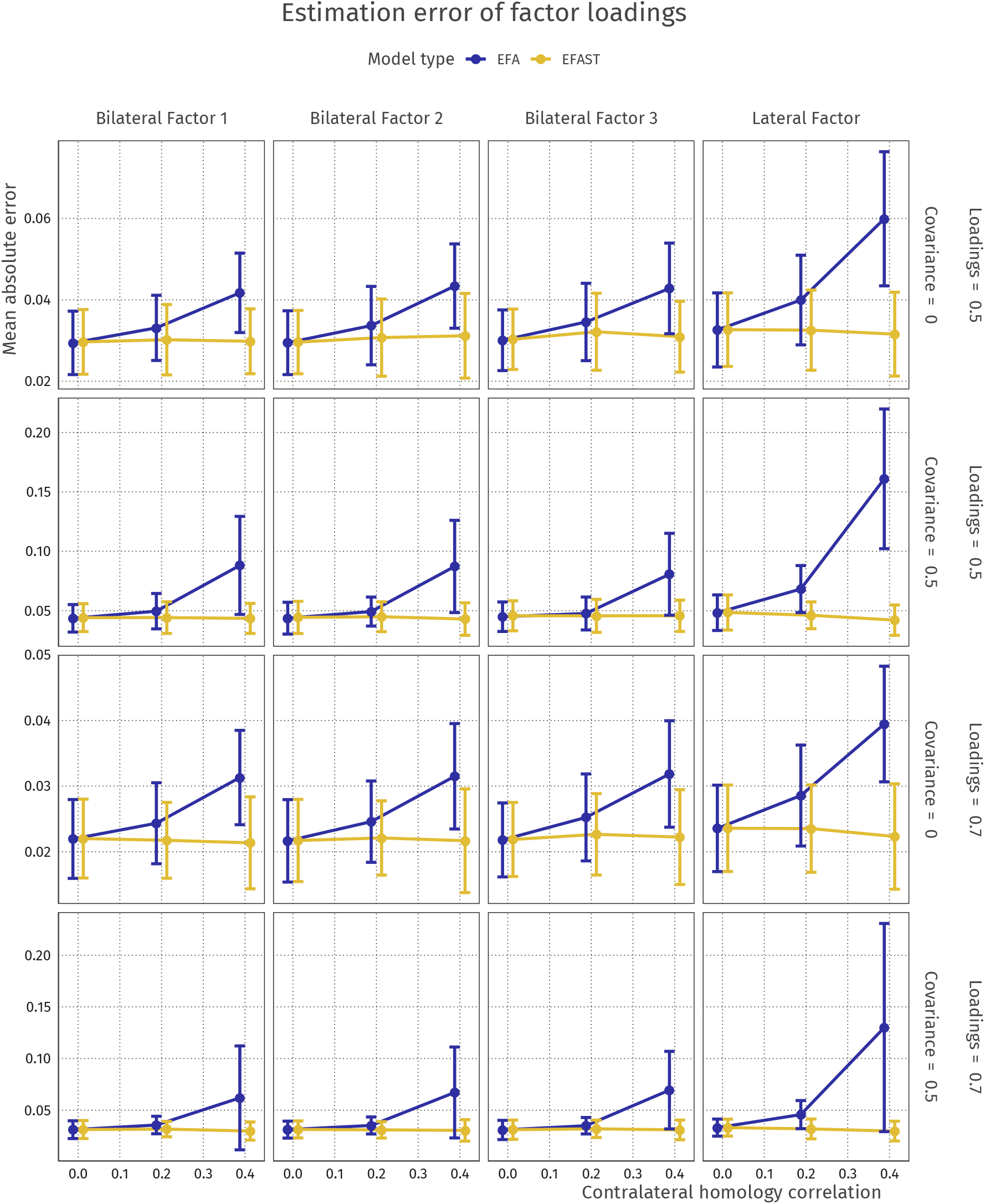
Factor loading median absolute error over different conditions of factor loading and factor correlation strength (top-to-bottom, see labels on the right) and different factors (left-to-right, see labels on top).

## C Information criterion factor extraction performance

**Figure 23:**
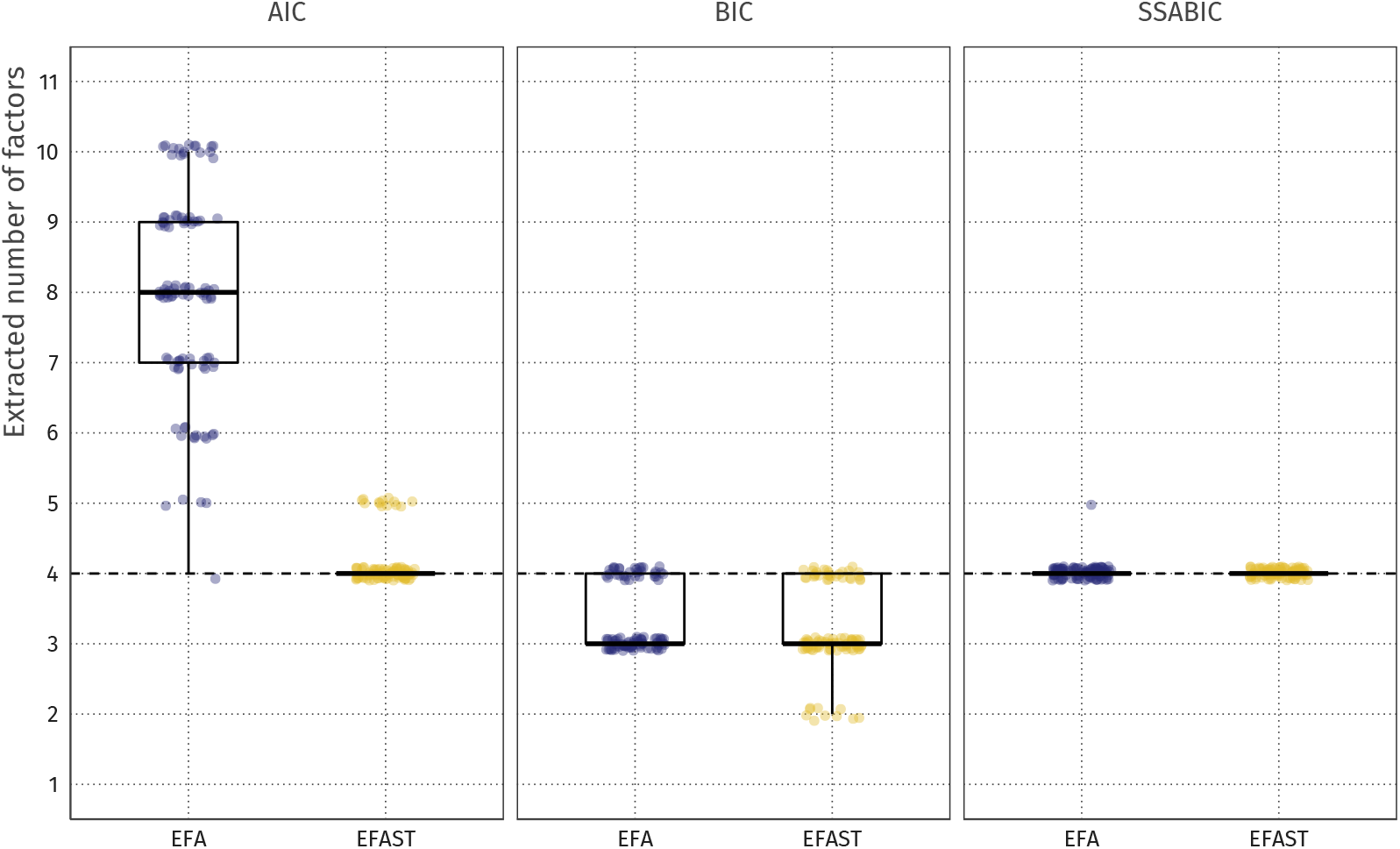
Number of extracted factors using the AIC (left panel), BIC (middle panel), and sample-size adjusted BIC (right panel) criterion. AIC works well for the EFAST method but not for the EFA method. BIC slightly underextracts for both methods. SSABIC shows excellent performance for both methods. The true number of factors is 4 (dashed line), for which this result holds; different simulation situations may show different factor extraction patterns.

## D Factor loadings for empirical application

**Table 4:** Factor loadings for 6-factor model fitted using EFAST and EFA on the Cam-CAN volume data. Loadings with an absolute value below 0.3 are not shown.

|  | EFAST |  |  |  |  |  | EFA |  |  |  |  |  |
| --- | --- | --- | --- | --- | --- | --- | --- | --- | --- | --- | --- | --- |
|  | F1 | F2 | F3 | F4 | F5 | F6 | F1 | F2 | F3 | F4 | F5 | F6 |
| lh_bankssts | 0.38 |  |  |  | 0.6 |  | 0.79 |  |  |  |  |  |
| lh_caudalanteriorcingulate |  |  |  | -0.4 | 0.55 |  |  |  |  |  |  |  |
| lh_caudalmiddlefrontal |  |  |  |  | 0.73 |  |  |  |  |  | 0.74 |  |
| lh_cuneus |  | 0.91 |  |  |  |  |  | 0.87 |  |  |  |  |
| lh_entorhinal |  |  | 0.31 |  |  |  |  |  |  |  |  |  |
| lh_fusiform |  |  |  |  | 0.4 |  | 0.38 |  |  |  |  |  |
| lh_inferiorparietal |  |  |  |  | 0.69 |  | 0.67 |  |  |  |  |  |
| lh_inferiortemporal |  |  |  |  | 0.4 |  | 0.54 |  |  |  |  |  |
| lh_isthmuscingulate |  |  |  |  | 0.44 |  |  |  |  |  |  |  |
| lh_lateraloccipital |  | 0.45 |  |  |  |  | 0.37 | 0.46 |  |  |  |  |
| lh_lateralorbitofrontal |  |  |  |  | 0.78 |  |  |  |  |  | 0.71 |  |
| lh_lingual |  | 0.71 |  |  |  |  |  | 0.7 |  |  |  |  |
| lh_medialorbitofrontal |  |  |  |  | 0.65 |  |  |  |  |  | 0.63 |  |
| lh_middletemporal | 0.31 |  |  |  | 0.66 |  | 0.81 |  |  |  |  |  |
| lh parahippocampal |  |  |  |  | 0.37 |  |  |  |  |  |  |  |
| lh_paracentral |  |  |  |  | 0.83 |  |  |  |  |  | 0.73 |  |
| lh_parsopercularis |  |  |  |  | 0.79 |  |  |  |  |  | 0.72 |  |
| lh_parsorbitalis |  |  |  |  | 0.61 |  |  |  |  |  | 0.63 |  |
| lh_parstriangularis |  |  |  |  | 0.84 |  |  |  |  |  | 0.89 |  |
| lh_pericalcarine |  | 0.91 |  |  |  |  |  | 0.91 |  |  |  |  |
| lh_postcentral |  |  |  |  | 0.8 |  | 0.37 |  |  |  | 0.33 |  |
| lh_posteriorcingulate |  |  |  |  | 0.72 |  |  |  |  |  | 0.52 |  |
| lh_precentral |  |  | -0.3 |  | 0.87 |  |  |  | -0.34 |  | 0.72 |  |
| lh_precuneus |  |  |  |  | 0.76 |  |  |  |  |  |  | 0.74 |
| lh_rostralanteriorcingulate |  |  |  |  | 0.59 |  |  |  |  |  | 0.49 |  |
| lh_rostralmiddlefrontal |  |  |  |  | 0.78 |  |  |  |  |  | 0.82 |  |
| lh_superiorfrontal |  |  |  |  | 0.88 |  |  |  |  |  | 0.93 |  |
| lh_superiorparietal |  |  |  |  | 0.72 |  |  |  |  |  |  | 0.77 |
| lh_superiortemporal |  |  |  |  | 0.77 |  | 0.52 |  |  |  | 0.41 |  |
| lh_supramarginal |  |  |  |  | 0.77 |  | 0.32 |  |  |  |  |  |
| lh_frontalpole |  |  |  |  | 0.32 |  |  |  |  |  | 0.51 |  |
| lh_temporalpole |  |  | 0.39 |  |  |  |  |  | 0.34 |  |  |  |
| lh_transversetemporal |  |  |  |  | 0.72 |  |  |  |  | 0.43 | 0.53 |  |
| lh_insula |  |  |  |  | 0.78 |  |  |  |  |  | 0.7 |  |
| rh_bankssts |  |  |  |  | 0.71 |  | 0.69 |  |  |  |  |  |
| rh_caudalanteriorcingulate |  |  |  | 0.73 |  |  |  |  |  |  | 0.48 |  |
| rh_caudalmiddlefrontal |  |  |  |  | 0.76 |  |  |  |  |  | 0.74 |  |
| rh_cuneus |  | 0.71 |  |  |  |  |  | 0.72 |  |  |  |  |
| rh_entorhinal |  |  | 0.34 |  |  |  |  |  | 0.34 |  |  |  |
| rh_fusiform |  |  |  |  | 0.5 |  | 0.4 |  |  |  |  |  |
| rh_inferiorparietal | 0.31 |  |  |  | 0.7 |  | 0.67 |  |  |  |  |  |
| rh_inferiortemporal | 0.31 |  |  |  | 0.38 |  | 0.58 |  |  |  |  |  |
| rh_isthmuscingulate |  |  |  |  | 0.45 |  |  |  |  |  |  |  |
| rh_lateraloccipital |  | 0.47 |  |  |  |  | 0.37 | 0.47 |  |  |  |  |
| rh_lateralorbitofrontal |  |  |  |  | 0.73 |  |  |  |  |  | 0.72 |  |
| rh_lingual |  | 0.66 |  |  |  |  |  | 0.68 |  |  |  |  |
| rh_medialorbitofrontal |  |  |  |  | 0.74 |  |  |  |  |  | 0.67 |  |
| rh_middletemporal |  |  |  |  | 0.66 |  | 0.75 |  |  |  |  |  |
| rh parahippocampal |  |  |  |  | 0.39 |  |  |  |  |  |  |  |
| rh_paracentral |  |  |  |  | 0.83 |  |  |  |  |  | 0.65 |  |
| rh_parsopercularis |  |  |  |  | 0.76 |  |  |  |  |  | 0.68 |  |
| rh_parsorbitalis |  |  |  |  | 0.62 |  |  |  |  |  | 0.82 |  |
| rh_parstriangularis |  |  |  |  | 0.71 |  |  |  |  |  | 0.8 |  |
| rh_pericalcarine |  | 0.77 |  |  |  |  |  | 0.82 |  |  |  |  |
| rh_postcentral |  |  |  |  | 0.77 |  | 0.31 |  |  |  | 0.35 |  |
| rh_posteriorcingulate |  |  |  |  | 0.57 |  |  |  |  |  | 0.44 |  |
| rh_precentral |  |  |  |  | 0.96 |  |  |  | -0.31 |  | 0.83 |  |
| rh_precuneus |  |  |  |  | 0.76 |  |  |  |  |  |  | 0.72 |
| rh_rostralanteriorcingulate |  |  |  | 0.44 |  |  |  |  |  |  | 0.51 |  |
| rh_rostralmiddlefrontal |  |  |  |  | 0.7 |  |  |  |  |  | 0.68 |  |
| rh_superiorfrontal |  |  |  |  | 0.95 |  |  |  |  |  | 0.81 |  |
| rh_superiorparietal |  |  |  |  | 0.82 |  |  |  |  |  |  | 0.71 |
| rh_superiortemporal |  |  |  |  | 0.91 |  | 0.39 |  |  |  | 0.46 |  |
| rh_supramarginal |  |  |  |  | 0.82 |  | 0.36 |  |  |  |  | 0.43 |
| rh_frontalpole |  |  |  |  |  |  |  |  |  |  |  |  |
| rh_temporalpole |  |  | 0.34 |  |  |  |  |  |  |  |  |  |
| rh_transversetemporal |  |  |  |  | 0.78 |  |  |  |  | 0.39 | 0.43 |  |
| rh_insula |  |  |  |  | 0.73 |  |  |  |  |  | 0.69 |  |

